# Cortical Hierarchy, Dual Counterstream Architecture and The Importance of Top-Down Generative Networks

**DOI:** 10.1101/2020.04.08.032706

**Authors:** Julien Vezoli, Loïc Magrou, Rainer Goebel, Xiao-Jing Wang, Kenneth Knoblauch, Martin Vinck, Henry Kennedy

## Abstract

Hierarchy is a major organizational principle of the cortex and underscores modern computational theories of cortical function. The local microcircuit amplifies long-distance inter-areal input, which show distance-dependent changes in their laminar profiles. Statistical modeling of these changes in laminar profiles demonstrates that inputs from multiple hierarchical levels to their target areas show remarkable consistency, allowing the construction of a cortical hierarchy based on a principle of hierarchical distance. The statistical modeling that is applied to structure can also be applied to laminar differences in the oscillatory coherence between areas thereby determining a functional hierarchy of the cortex. Close examination of the anatomy of inter-areal connectivity reveals a dual counterstream architecture with well-defined distance-dependent feedback and feedforward pathways in both the supra- and infragranular layers, suggesting a multiplicity of feedback pathways with well-defined functional properties. These findings are consistent with feedback connections providing a generative network involved in a wide range of cognitive functions. A dynamical model constrained by connectivity data shed insights into the experimentally observed signatures of frequency-dependent Granger causality for feedforward versus feedback signaling. Concerted experiments capitalizing on recent technical advances and combining tract-tracing, high-resolution fMRI, optogenetics and mathematical modeling hold the promise of a much improved understanding of lamina-constrained mechanisms of neural computation and cognition. However, because inter-areal interactions involve cortical layers that have been the target of important evolutionary changes in the primate lineage, these investigations will need to include human and non-human primates comparisons.

**Plan:**

1. Introduction
2. Hierarchy – signatures of inputs to the local circuits.
3. Models of hierarchy
4. Hierarchy – input consistency
5. Dual stream architecture.
6. Functional characteristics of FF and FB pathways.
7. The predictive brain and the importance of top-down generative networks.
8. Conclusion.

## 1. Introduction

Hierarchy as an organizational feature of the brain has been a recurrent theme since the evolutionary theory of neurological disorders of John Hughlings Jackson (1835-1911), following his observations of positive and negative symptoms in his patients (York and Steinberg, 2011). The neurobiology of cortical hierarchy was explored by the pioneering work of David Hubel and Torsten Weisel when they characterized the receptive field properties of simple, complex and hypercomplex neurons across areas of the visual cortex (Hubel and Wiesel, 1962). Following the work of Rockland and Pandya (1979) a myriad of connectivity studies in the cortex found additional evidence of hierarchical organization, allowing Felleman and Van Essen to propose the first hierarchical model of the cortex (Felleman and Van Essen, 1991), thereby providing a framework for modern concepts of feedforward (FF) and feedback (FB) processes. The notion of hierarchy has become considerably generalized and for example can be considered to be the major concept linking biological and artificial intelligence (Hawkins and Blakeslee, 2004). Convolutional deep neural networks have a clear hierarchical organization, with convergent, FF connections passing information from lower to higher layers, and divergent FB connections shaping plasticity in the connections from lower layers (LeCun et al., 2015). But what exactly is the significance of hierarchy in the brain? Hierarchy has been extensively studied in terms of ranking of cortical areas with respect to a number of criteria including, gradients of structural and functional features, as a progression of scales or as a topological sequence of projections (Hilgetag and Goulas, 2020). Here we take a diametrically opposing view. Rather than simply considering hierarchy as a ranking of cortical areas, we address what it means in terms of monosynaptic inter-areal connectivity. In line with the tenet that the explanation of how the brain works demands an account of what neurons do, and that functional interactions of cortical areas is assured by neuronal activity relayed between areas by axons, we confine our discussion of hierarchy to the description of the neuronal properties of inter-areal relations. We propose that the structural and functional markers of hierarchy define the integration of long-range inputs into each local circuit. Future investigation of these markers are expected to provide insight to the cellular mechanisms underlying hierarchical processing. A critical aspect of these regularities concerns the spatial distances governing interactions between cortical neurons, which we and others have shown obey metabolic constraints in terms of wire minimization underlining the spatial embedding of the cortex (Markov et al., 2013). We discuss the functional implications of the Dual Stream Architecture (Markov et al., 2014b) in terms of FF and FB processing and where future investigations are expected to provide insight into the cellular mechanisms underlying hierarchical processing. Finally, we cast FF and FB relations in terms of predictive processing theory and evoke the importance of top down generative networks.

## 2. Hierarchy – signatures of inputs to local circuits

In 1989 Douglas, Martin and Whitteridge published a landmark study that proposed a canonical microcircuit for the neocortex (Douglas et al., 1989) (**Figure 1 and 10b**). A common misconception of the canonical microcircuit is that it constitutes solely a description of the inter-laminar wiring patterns of the cortex. In fact, it is much more a theory that sets out to explain the electrophysiological properties of the cortex in terms of local connectivity, in particular with respect to within-laminar connections. In an effort to nail down the transformation of the thalamic input, *in vivo* intracellular recordings were made in area V1 in cat cortex. This showed that minute inputs from the LGN are amplified by recurrent excitation in layer 4 neurons (Latawiec et al., 2000). Subsequent quantification of the synaptic components of the local circuit showed that high levels of within-layer recurrent excitation is a characteristic feature of the local circuit (**Figure 1**) (Binzegger et al., 2009). These experiments showed that the role of inhibition was not to carve out the selectivity of the neuron response but rather to exert a control over the amplitude of the response and therefore to maximize the inhibitory potentials in the optimal receptive field response (Douglas et al., 1995; Douglas et al., 1989). Subsequent work showed that there is a weak orientation response in the input to the cortex, meaning that the primary role of the recurrent excitation is the amplification of a signal and not its creation (Ferster et al., 1996).

**Figure 1.**
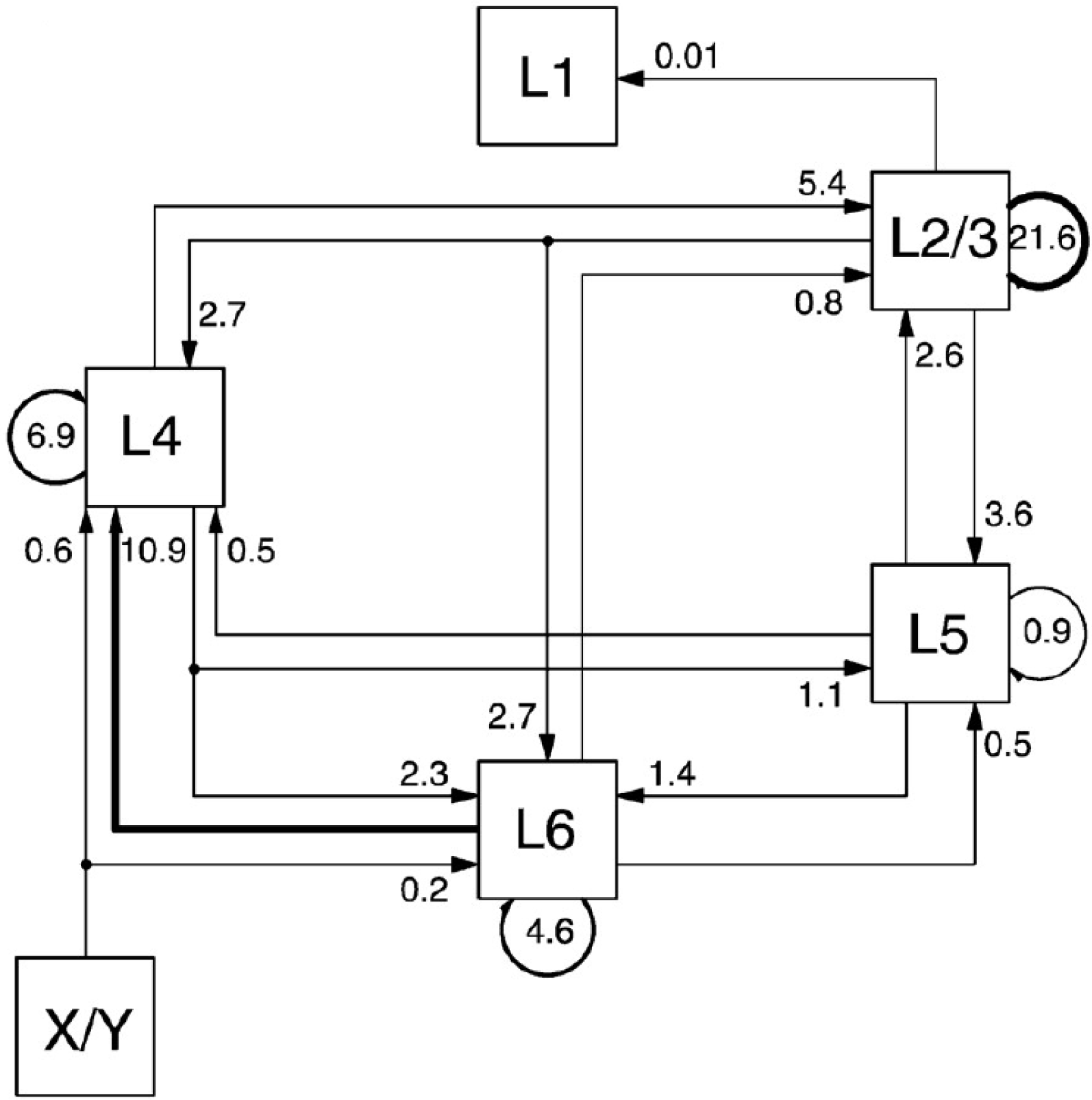
Quantitative map of excitatory synapses between excitatory neurons of the local microcircuit in visual cortex (area 17) of the cat. Numbers indicate proportions of excitatory synapses, note the dominance of within layer recurrent connectivity with 21.6 peak values in Layers 2/3. The FF loop starts in layer 4, the major thalamic recipient layer and then extends to layers 2/3, 5 and 6 with recurrent inputs back to layer 4. This FF loop corresponds to a little less than half of synapses involved in self-innervation of individual cortical layers. X/Y refers to the component cells of the lateral geniculate nucleus, the major thalamic relay. The original canonical microcircuit is shown in Figure 9B. L refers to layer. From (Binzegger et al., 2004) with permission.

For many years research on cortex was predominantly in carnivores and non-human primates, leading to the notion of the cortical column as a fundamental component of functional organization (Mountcastle, 1995). In these studies, electrophysiological recordings from electrode penetrations made perpendicular to the cortex found a conserved function in the depth of the cortex in passing from pia to white matter (Hubel and Wiesel, 1962; Mountcastle, 1957). In the visual system there were expectations that the columnar organization of the cortex would be both functional and structural, since physiologically demonstrated ocular-dominance columns appeared to co-localize with cortical territories labeled by transynaptic labeling following tracer injections in the eye (Hubel and Wiesel, 1977). However, close examination revealed important discrepancies in such a correspondence (da Costa and Martin, 2010)), suggesting that the link between cortical structure and function is to be found at a much finer scale. Thanks to work in the mouse visual cortex using the molecular tools that are available in this species, it has been confirmed that cortical responses to thalamic input are indeed the consequence of an amplification (Harris and Mrsic-Flogel, 2013) (Lien and Scanziani, 2013) via the local recurrent excitation (Cossell et al., 2015; Douglas et al., 1995; Ko et al., 2011). These studies built on earlier findings of highly nonrandom features of synaptic connectivity in local cortical circuits, proposing that there is a skeleton of strong connections in a sea of weak connections (Song et al., 2005). Later it was shown that the rare strong connections in the lognormal distribution of synaptic weights are between neurons with similar receptive fields, meaning that neurons in the visual cortex listen most closely to a subset of their synaptic inputs (Cossell et al., 2015). These findings explain earlier observations showing that ongoing activity of a neuron (so called spontaneous activity) reflects the functional architecture (i.e. the backbone of strong connections) in which it is embedded (Tsodyks et al., 1999). The emerging picture is that layers 4, 3 and 2 neurons are organized into subnetworks so that the selectivity of the amplification is ensured by constraints at the scale of dendritic spines (Lee et al., 2016). The principal wiring property of the canonical circuit is the recurrent excitation that is observed in all of the cortical layers including layer 4 (Binzegger et al., 2004). The relevance of the canonical microcircuit theory for understanding inter-areal processing became apparent when cortical connectivity was quantified. In fact, 80-90% of the connections of the cortex are in the local circuit spanning 1-2mm in the visual cortex (Markov et al., 2011). Except for the adjacent cortical area, the structural weight of the average input from a distant source area to a target area is several orders of magnitude *less* than the thalamic input (Markov et al., 2014a). These observations lead to the notion that amplification by local recurrent excitation is a general phenomenon, that allows selection and recombination of relatively small afferent signals (Douglas and Martin, 2007a, b). For instance, top-down signaling of selective attention multiplicatively modulates sets of sensory neurons (McAdams and Maunsell, 1999; Treue and Maunsell, 1996). In this manner, selective amplification by local circuit dynamics leads to all-or-none task switching (Ardid and Wang, 2013).

Early anatomists, working principally in non-human primates, distinguished between rostral directed connections that originate chiefly in the supragranular layers and terminate in layer 4 (Cragg, 1969; Kennedy and Bullier, 1985; Lund et al., 1975; Martinez-Millan and Hollander, 1975; Rockland and Pandya, 1979; Spatz et al., 1970; Van Essen and Zeki, 1978; Wong-Riley, 1978) and caudal directed connections that mostly originate from infragranular layers and terminate outside of layer 4 (Kaas and Lin, 1977; Kennedy and Bullier, 1985; Kuypers et al., 1965; Tigges et al., 1973; Wong-Riley, 1978). In a landmark study, Rockland and Pandya (Rockland and Pandya, 1979) were the first to formulate inter-areal connectivity in terms of hierarchy and suggested that the laminar organization of cortical afferents and their terminals indicates the sequence of information processing in cortex. These authors proposed that connections originating from supragranular layers and terminating in layer 4 by analogy with the main thalamic input to cortex constitute the FF pathway channeling sensory information to cortical areas carrying out higher-order analyses. By contrast connections arising from the infragranular layers, by analogy with descending projections to subcortical structures, correspond to FB connections and were postulated to enable higher order areas to be able to modulate the activity of lower level areas (Rockland and Pandya, 1979).

## 3. Models of Hierarchy

The classification of pathways between areas as FF and FB helped motivate the investigation of the role of the cortical network in terms of FF pathways shaping receptive fields in their target areas (Hubel, 1995) and FB pathways relaying contextual information (Gilbert and Li, 2013; Zipser et al., 1996). How the cortical network related to the mosaic of cortical areas was given substance by Felleman and Van Essen’s demonstration that the layout of cortical areas corresponded to a distributed hierarchy (Felleman and Van Essen, 1991). In their seminal study these authors established a definition of FF and FB connections largely employing the criteria of Rockland and Pandya (1979), and although principally based on laminar patterns of anterograde labeling they were able to stipulate criteria so as to include retrograde labeling therefore enabling them to define pathways with respect to findings reported in a large number of publications (**Figure 2A**). Pairwise comparisons of the connections linking areas using these criteria revealed a high regularity; connections that were classified as FF were largely reciprocated by FB connections, allowing the authors to establish a distributed hierarchy across multiple streams in the macaque visual cortex shown in **Figure 2B**. Because of the many parallel pathways and given that hierarchical levels were defined arbitrarily, it meant that the precise ordering of cortical areas was ill-defined. Computational modeling showed that there were over 150,000 equally plausible solutions to the Felleman and Van Essen Model (Hilgetag et al., 1996) (**Figure 2C**).

**Figure 2.**
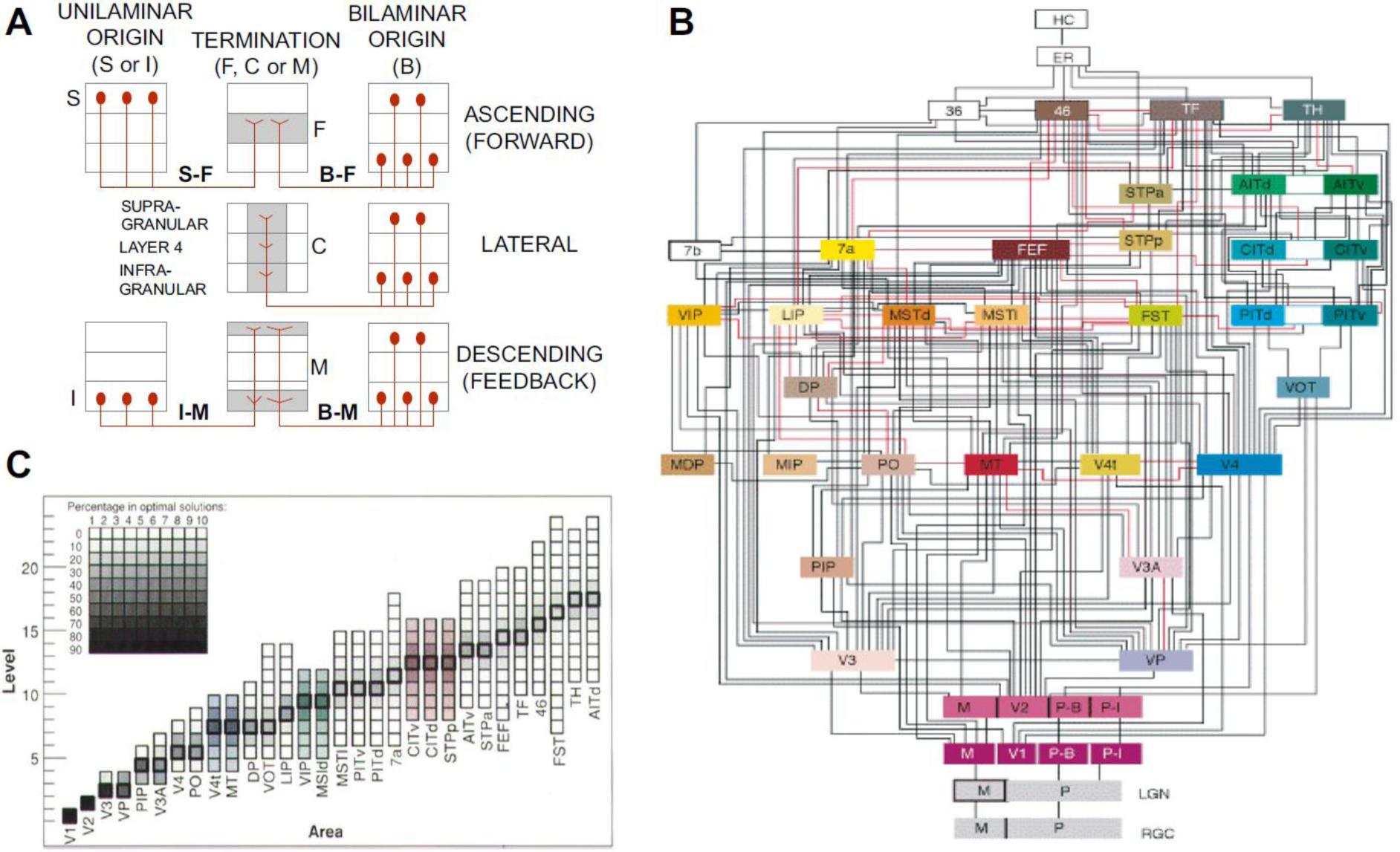
The Felleman and Van Essen binary model of cortical hierarchy. **A)** Criteria for classifying connections between areas as FF (top), lateral (middle) and FB (bottom) row. Termination patterns are depicted in the central column, preferentially in layer 4 (F pattern) FF, across all layers (C pattern) lateral, in upper and lower layers avoiding layer 4 (M pattern) FB. Laminar origin from a single layer (left column), is either supragranular (S) and therefore FF, or infragranular (I) and therefore FB. Bilaminar (B) origins (right column) either terminate in the middle layers (F pattern) and are therefore FF, terminate in all layers (lateral) or terminate predominantly in upper supra- and infragranular layers (M pattern) and therefore FB. **B)** The binary hierarchical model. **C**: Area frequency distributions for 150,000 optimal hierarchical orderings (Hilgetag et al., 1996).

A solution to the indeterminacy of the Felleman and Van Essen model could be overcome by an objective localization of hierarchical level. A suggestion that this might be the case was the observation that injections of retrograde tracers in a target area in one of the early visual areas generated a progressive decrease in the numbers of labeled FB neurons in supragranular layers in source areas with increasing physical rostral distance (**Figures 3A**) (Kennedy and Bullier, 1985). Quantitative measures of interareal connectivity showed that patterns of retrograde labeling were highly consistent across different brains, provided that labeled neurons are summed across the full extent of a projection zone, defined as the region in a particular source area which contains projections to an injected target area (**Figure 3B,C**). Subsequently injections in cortical areas at higher hierarchical levels generated a progressive increase in the numbers of labeled FB neurons in supragranular layers with increasing physical caudal distance. In this manner FF and FB pathways exhibited opposing gradients of projection neurons (Barone et al., 2000; Kennedy and Bullier, 1985; Markov et al., 2014b). These observations led to the definition of an index of this gradient based on the proportion of Supragranular Labelled Neurons or SLN (Barone et al., 2000; Vezoli et al., 2004). Because these changes are highly consistent across brains, the smooth gradients of inputs from neurons in different layers and areas to a target area lead to the derivation of a hierarchical distance rule (**Figure 3B**).

**Figure 3.**
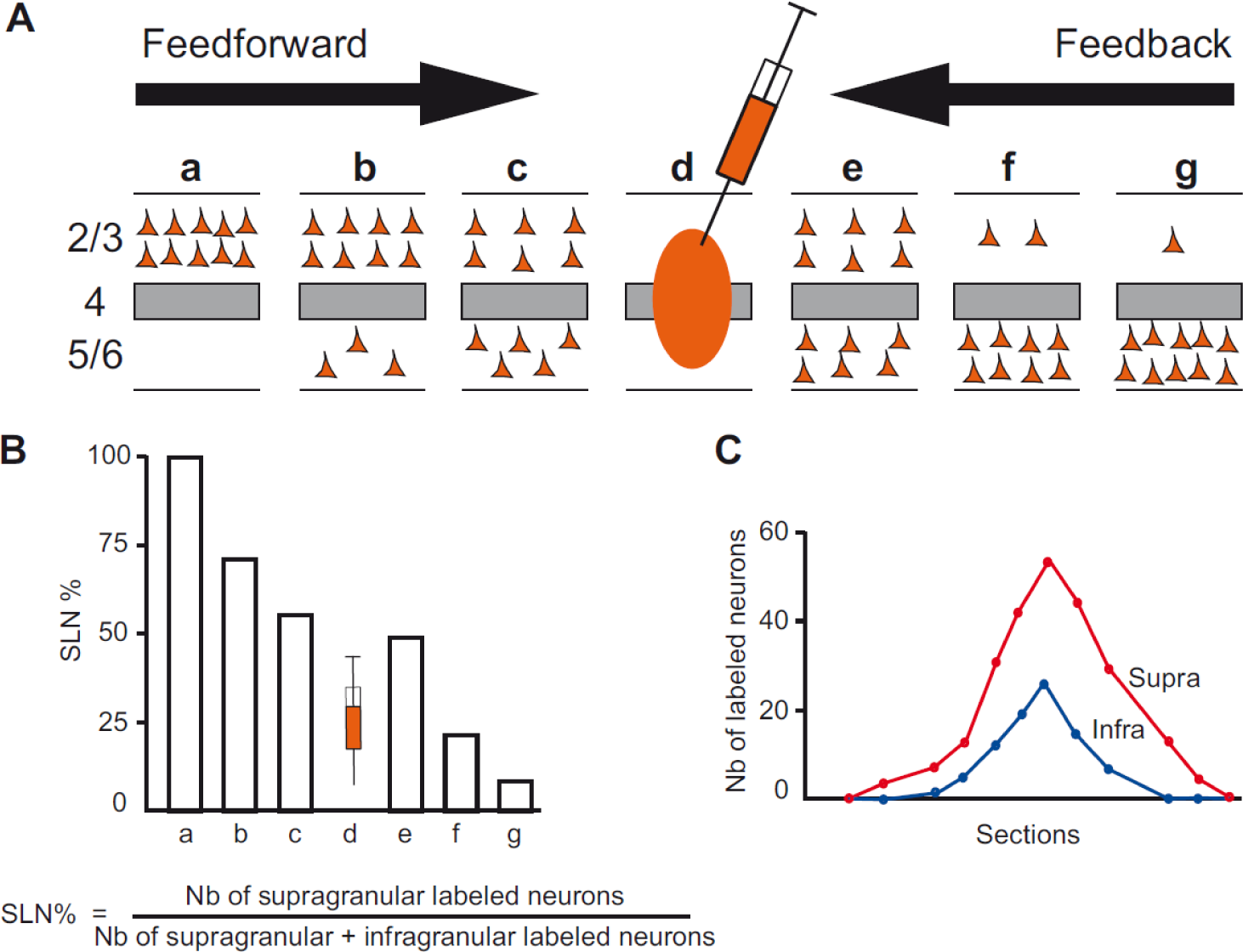
Quantitative parameters characterizing the hierarchy. **A**) The laminar distribution of parent neurons in each pathway, referred to as SLN (fraction of supragranular neurons) is determined by high frequency sampling and quantitative analysis of labeling. Supra- and infragranular layer neurons contribute to both FF and FB pathways, and their relative proportion is characteristic for each type of pathway. For a given injection there is a gradient of SLN of the labeled areas, between purely FF (SLN = 100%, all the parent neurons are in the supragranular layers) to purely FB (SLN = 0%, all the parent neurons in the infragranular layers) and a spectrum of intermediate proportions; **B**) All labeled areas can then be ordered by decreasing SLN values and this order is consistent with hierarchical order according to Felleman and Van Essen. SLN is thus used as an indicator of hierarchical distance between areas from the same injection; **C**) Reliable estimation of SLN crucially requires sampling labeling throughout the full extent of the projection zone in each area.

The transition from a binary model of hierarchy to one based on hierarchical distance had important consequences. One way of thinking about these connectivity gradients (**Figure 3A**) is that they represent gradual changes in the composition of inputs to the local microcircuit of a cortical area that is dependent on physical distance. Elsewhere we have shown that input strength to cortical areas declines exponential with distance (Ercsey-Ravasz et al., 2013). In terms of hierarchy, the strong nearby connections are near-lateral and with increasing distance connections are progressively more FB or FF in nature.

If these changing inputs to the local microcircuit represent the structural signature of hierarchy it is legitimate to wonder if they have a functional correlate? If this is the case, then Pascal Fries and his team reasoned that one can derive a functional hierarchy (Bastos et al., 2015b). The hierarchical distance rule is based on the fact that supragranular layers primarily send FF projections and infragrananular layers FB projections. In the visual system, superficial and deep layers are characterized by relatively strong gamma and alpha/beta oscillations, respectively (Buffalo et al., 2011). Furthermore, whereas in early visual areas, gamma oscillations are relatively strong (Gray et al., 1989), beta oscillations tend to be strong in higher areas like those in parietal cortex (Brovelli et al., 2004; Scherberger et al., 2005). These observations lead to the prediction that in the visuo-parietal system interareal synchronization in the gamma frequency band mediates FF and interareal synchronization in the alpha- and beta-frequency band mediate FB influences. Granger-causality was used to quantify functional FF and FB interactions, thereby allowing the strength of these interactions to be quantified for individual frequency bands. Neuronal populations show spontaneous fluctuations over time that are driven by brain rhythms in different frequency bands, such as theta (3-8Hz), alpha (8-12Hz), beta (12-30Hz) and gamma (30-80Hz). Note, neocortical rhythms do not comprise band-limited sinusoidal oscillation but typically describes noisy oscillations with energy in a wide frequency range (Burns et al., 2011; Spyropoulos et al., 2020). Frequency-specific directed influences of rhythmic synchronization are correlated with hierarchical distance, FF pathways are characterized by synchronization in the theta (4 Hz) and gamma-band (60-80 Hz) and FB in the beta-band (14-18 Hz) (Bastos et al., 2015b). These observations mean that the structural signatures to the microcircuit are indeed paralleled by functional signatures (Bastos et al., 2015b). However, whereas the structural hierarchy is fixed theses authors were able to show that activity patterns underlying functional hierarchy exhibit task dependent dynamics.

Using empirical connectivity data as a structural substrate, Mejias et al. (Mejias et al., 2016) built a dynamical model of multi-regional macaque monkey cortex endowed with a laminar structure. The model captures stochastic fast oscillations in the gamma frequency range in the superficial layers, and lower-frequency (alpha) oscillations in the deep layers. Importantly, in contrast to regular oscillators, such network rhythms are quite irregular and noisy, compatible with the notion of sparsely synchronous brain rhythms (Wang, 2010), which provides a unifying framework for explaining both population oscillations and Poisson-like highly variable spike trains of single neurons. The model accounts for the distinct signatures of frequency-dependent Granger causality that reflect FF versus FB signaling, and reproduces the experimentally deduced neural population dynamics that are consistent with the anatomically defined hierarchy. Therefore, this model provides a computational platform for theoretical investigations of the interplay between feedforward and feedback processes.

Both structural and functional hierarchies show that the regularities stemming from laminar distributions of connections and the signals they confer are remarkably consistent. In the following section, we address the extent of this consistency in order to formalize how hierarchy inputs to an area are shaped by distance.

## 4. Hierarchy – Input Consistency

The notion of hierarchy implies order or rank, so that a prerequisite to determining if there are hierarchical relations between cortical areas requires determining if order relations can be defined between them. For example, the Felleman and Van Essen hierarchy was based on the binary classification of FB/FF relations between areas defined by laminar origin and termination of projections (Felleman and Van Essen, 1991). A FF projection from area A to B implied that B was ordered after A. Similarly, a FB projection from B to A would also be consistent with the above FF relation in assigning B after A. While in a hierarchy we would expect the two criteria to agree that might not in fact be the case. On a simple level, disagreement could be taken to define equality of ordinal levels in the sense that equality is defined as those numbers, A and B, that simultaneously satisfy *A*≥*B* and *A*≤*B*. Alternatively, distinct hierarchical orders might arise: one on the basis of FF laminar projections, the other on the basis of FB. These observations become important when the data supporting multiple laminar FB/FF pathways are analyzed.

The criteria for determining hierarchical rank described above are based on the properties of projections that define relations between areas. Alternatively, one can consider properties that are intrinsic to an area, such as cortical thickness, neuron density, soma size, spine count, spine density and dendritic tree size. These properties have been shown to display a smooth variation across the cortex that allows ranking of areas in accordance with the gradation principle of Sanides (Barbas, 2015; Sanides, 1972). Because these properties vary across the cortex, a hierarchical ranking can be established simply by ordering the areas according to the property. This distinction leads us to consider that criteria for building cortical hierarchies can be divided into two broad classes that we shall refer to as *node-based* and *link-based* (**Figure 4A**).

**Figure 4.**
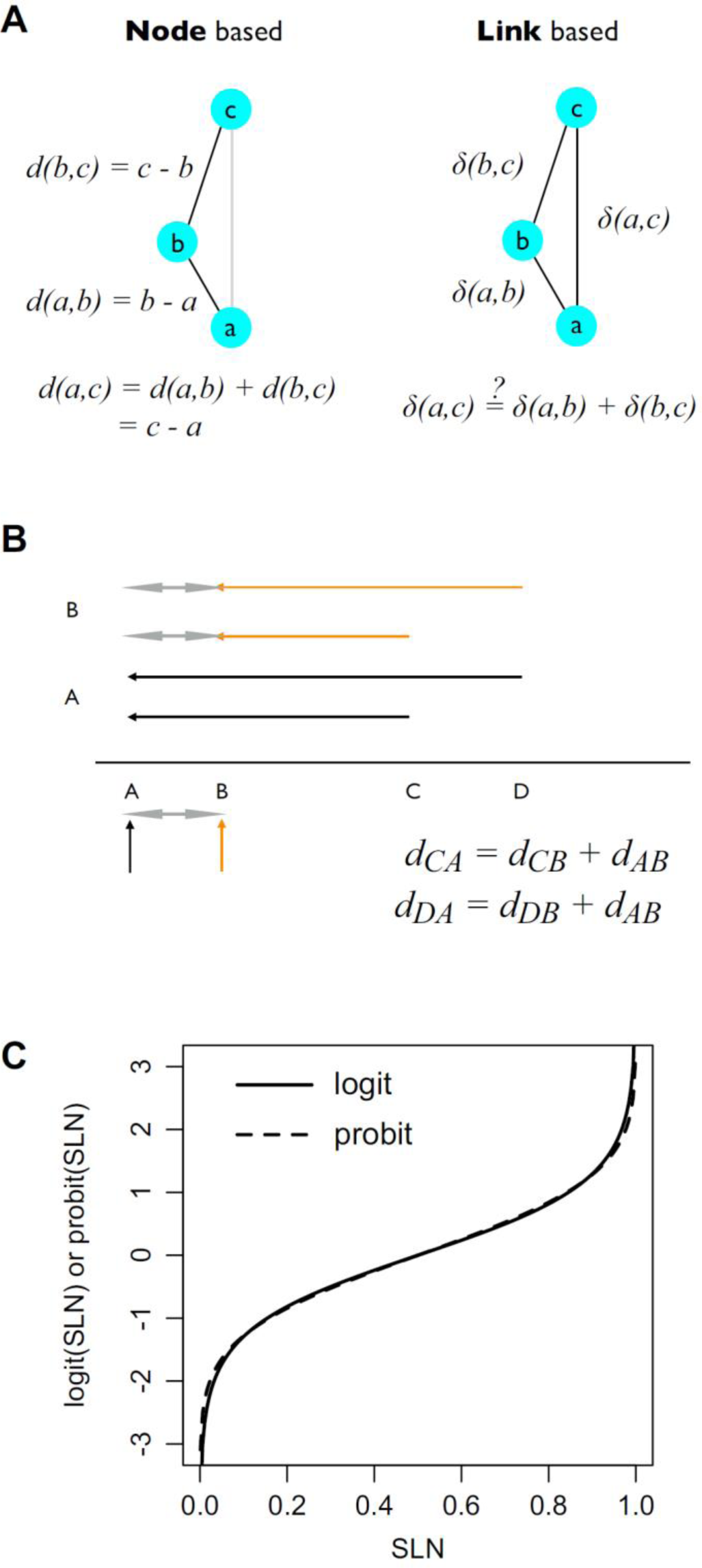
Properties of nodes and links. Nodes have fixed properties, a, b, c which in turn fixes their order and any distance measure, d, calculated from these properties. b. Link properties depend on the relations between node pairs, ab, bc, ac. The distance measures, δ, for ab, and bc do not necessarily fix that for ac. The above graphs are unidirectional, but in bi-directional graphs the distances between nodes need not by symmetric. **B) Hierarchical scales**. Suppose a hierarchical scale between areas A, B, C, D, with the ordering and distances as illustrated on the bottom line. We expect measures of distance to be consistent measured between any pairs of areas. For example, injections in areas A and B lead to distances defined with respect to each of these areas, i.e., distances AB (double headed grey arrow), AC and AD (black arrows) for injection in area A, and BA (double headed grey arrow), BC and BD (orange arrows) for injection in area B. Consistency would imply, for example, that for a distance measure, d, the estimate of d_AB_ = d_BA_ would be the same for both injections, i.e., d_CA_ – d_CB_ = d_DA_ – d_DB_. **C) SLN Transformation**. Comparison of logit (solid) and probit (dashed) transformations of SLN values on the interval (0, 1). The logit SLN is defined as ln(SLN/(1 – SLN)). The probit is defined as the inverse of the Gaussian cumulative distribution function and is often notated by Φ^−1^. The scale factor of the logit curve has been adjusted by a factor of 0.588 to match it to the probit curve.

Here it is useful to draw an analogy with social networks. A hierarchy in a social network implies that the actors show specific kinds of interactions with each other (*link-based*). Hierarchy implies that those close to each other in a hierarchy show stronger interactions with each other than actors that are distant in the hierarchy. More information can be gauged from the nature of the interactions: We expect that someone high in the hierarchy (a boss) will show a more directive communication pattern to workers lower in the hierarchy. The workers, in turn, will show a different ascending communication pattern, e.g. asking more questions. Thus, a hierarchy can be constructed by studying the way in which people interact with each other, and knowing a hierarchy could in principal allow us to predict these interactions. By analogy, the SLN can be seen as a measure that directly quantifies the nature of the interactions between brain areas, based on the laminar projection patterns. Interestingly and as we described above, these laminar projection patterns also relate to certain functional interaction patterns (FF gamma and FB alpha/beta influences). In addition, social hierarchy might also be gauged from properties of the people in the hierarchy themselves. For instance, one expects the boss of the company to have the largest office, whereas the workers at the bottom to share an office and have smaller desks (*node-based*). In some sense, one could argue however, that the *node-based* view is based only on indirect markers and is ultimately grounded in the interaction *link-based* view.

There are critically important differences for constructing hierarchies between node and link-based information. By definition, node-based relations depend only on a value intrinsic to the node, not the relations between nodes so they give no information on the symmetry or otherwise of inter-areal relations. By contrast, ranks based on links are expected to show reciprocity, so that if there is FF pattern from area A to area B, a FB pattern is expected from area B to area A. Node based criteria are defined between *any* two areas independently of whether or not a connection is actually present. Link-based criteria can provide information on asymmetric relations, provided they are directional and are strictly defined between areas *only* when there is a direct connection. Nevertheless, hierarchical ordering between unconnected areas can be achieved through indirect connections. Generally, link-based hierarchy describes the connections that are carrying information between areas and therefore the manner in which the connections and activity from source areas are integrated into the local circuit of the target area.

In order to define a hierarchical distance scale, i.e., that is not just ordinal, a distance function, *d* has to be defined. This function *d* should transform whatever anatomical or physiological measure one is using into a consistent measure of hierarchical distance across cortical areas. For example, **Figure 4B** shows a hypothetical distance scale on which 4 areas, A, B, C, D, are arranged. Suppose that hierarchical distances are estimated based on measures derived from tracer injections in areas A and B. The injection in area A provides information about hierarchical distances to areas B, C and D and the injection in B to areas A, C and D. A consistent measure of hierarchical distance, *d*, would generate the same estimate of distance, *d*_AB_, between areas A and B or, formally, we would expect that *d*_CA_ *– d*_C*B*_ *= d*_DA_ *– d*_D*B*_. This is easily derived from the two equations in **Figure 4B** that show for two areas, C and D, the expected relation between the hierarchical distances of a common area to C or D to two areas sites, A and B. For common projections X, plotting *d*_XA_ against *d*_XB_ should fall along a line of unit slope with intercept *d*_AB_. The question is how to define the distance function *d*.

In contrast to the binary measure of hierarchy in the Felleman and Van Essen model, SLN is a continuous measure on the interval (0, 1), thus providing a measure of the degree of feedforwardness/feedbackness. A binary description treats a projection as FF (FB) if its SLN is greater (less) than 0.5. Using simply the SLN differences as a hierarchical distance measure, the Barone et al., 2000 study was able to reproduce nearly all of the features of the Felleman and Van Essen model based on the SLN values from injections in just two areas, V1 and V4. A notable exception is that the Barone et al. 2000 study placed the frontal eye field (FEF), this prefrontal area, *below* the early extrastriate visual area V4. The SLN value from FEF to V4 was above 0.7 which placed V4 at higher hierarchical levels. Subsequent physiological studies confirmed an FEF role in early visual processing (Moore and Armstrong, 2003; Schall, 2015), thus justifying its relatively low hierarchical level. The unusual FF pattern for such a caudally directed projection was further confirmed in other studies (Pouget et al., 2009) and pertains to a specific link-category on which we expand later.

While differences in SLN establish a determinate hierarchical distance measure between areas, the measure is not necessarily consistent in the manner described in **Figure 4B**. As the measure is defined on the interval (0, 1), SLN differences for two areas projecting to a third area could be quite different from those to another more distant area. An ideal measure would project the interval (0, 1) to a scale where differences remain linear. This is commonly accomplished in statistical models, such as generalized linear models (GLM), by means of a logit or probit transformation (**Figure 4C**) that map the unit interval onto the real line. As the figure demonstrates, with the proper scaling both of these transformations yield rather similar mappings.

**Figure 5** shows a set of scatter plots for SLN values of common projections for all area pairs between each of 11 visual areas injected with retrograde tracer (Markov et al., 2014b). The SLN values are plotted on probit transformed axes. For many of the area pairs, the point distributions are reasonably well described by a line of unit slope (dashed blue in each graph), as predicted by a consistent measure of distance, i.e., g(SLN_j_) = g(SLN_i_)+ c, where c is a constant. Given the similarity of the transforms, it is not surprising that the logit transformation yields virtually the same linear patterns between area pairs. Thus, this indicates that the ratio of supra- and infra-granular projections follows a gradient across the cortical areas and constitutes a global signature of hierarchical distance among these areas.

**Figure 5.**
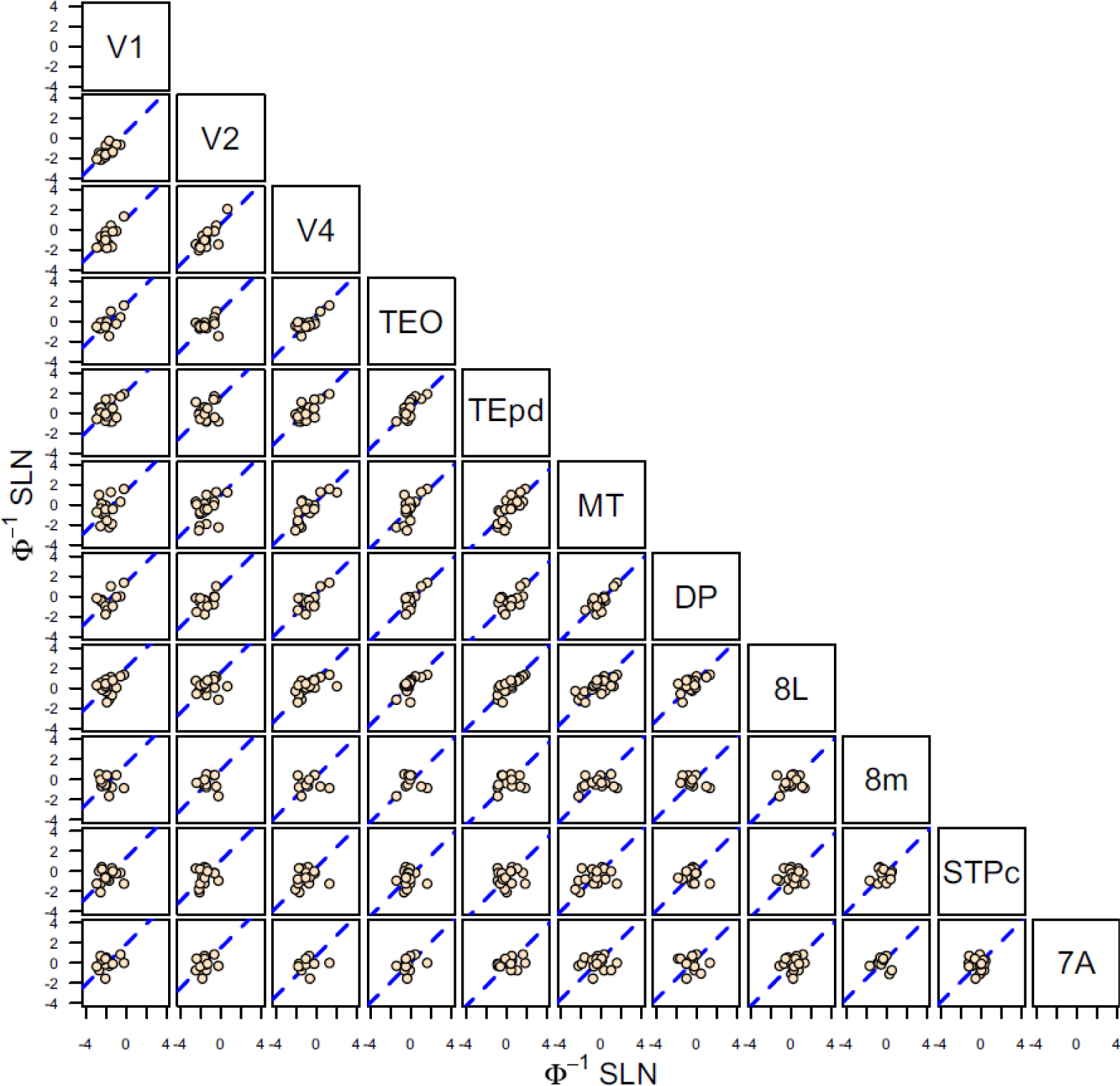
Probit transformation. Scatter plots of probit transformed SLN values of common source areas from pairs of 11 visual areas, obtained from retrograde tracer injections. The absicssa of each graph corresponds to the transformed SLN values of area *i*, indicated on the diagonal at the top of the column and the ordinate values are the transformed SLN values of area *j* indicated on the diagonal at the right of the row. The dashed blue line in each plot is the best fit line of unit slope (replotted from (Markov et al., 2014b)).

Is this laminar pattern of connectivity specific to the visual system or is it a more general principle of organization of FF and FB pathways in the brain? In support of the latter hypothesis, **Figure 6** shows paired scatter plots of SLN values for a set of 8 somatosensory and motor areas in the same format as **Figure 5**. As in the visual system, the transformed SLN values, here by the similar logit function, provide evidence of a consistent distance measure in the hierarchical organization among these areas.

**Figure 6.**
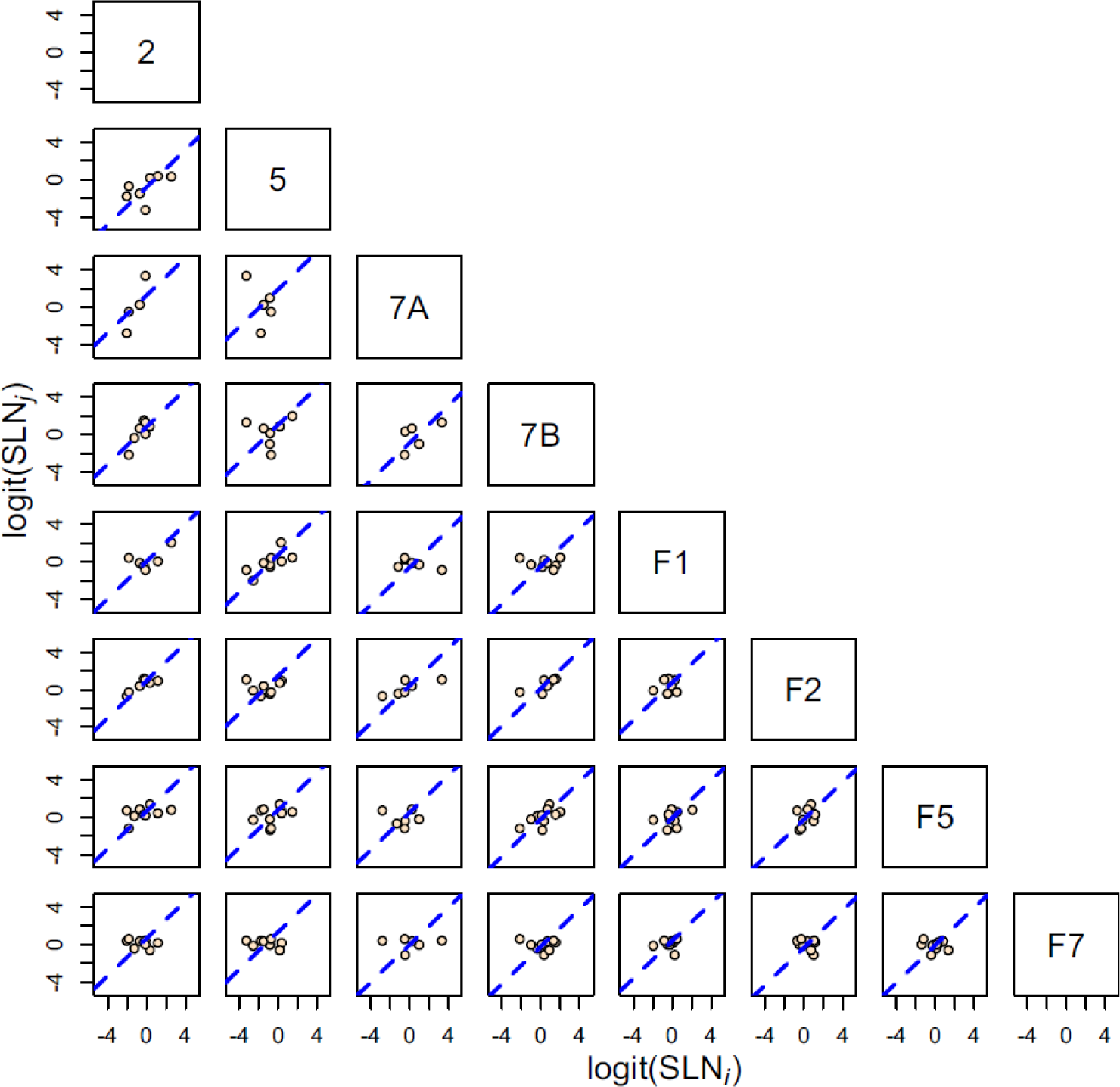
Logit transformation. Scatter plots of logit transformed SLN values of common source areas from pairs of 8 somatosensory and motor areas, obtained from retrograde tracer injections. The plots follow the same format as in Figure 6 except that the SLN values from each axis are transformed by the logit function. The dashed blue line in each plot is the best fit line of unit slope.

To quantify the consistency displayed in these pairs plots, we proposed a model to estimate hierarchical distances based on SLN values, but as we argue below, the model is quite general in its application. In short, we suppose that we can assign hierarchical levels, *h*_*i*_ and *h*_*j*_, to all area pairs *i* and *j*, based on a measure of differences between properties linking the areas. For example, in the case of SLN, we suppose

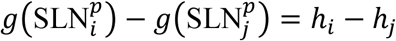

where *g* applies a logit or probit transformation to SLN values, from an injection into area *p* that receives projections from areas *i* and *j*. This suggests a formalism similar to a GLM with a binomial family. The SLN is taken as a binomial variable (neurons are found in the upper or lower cortical layers) and the sum of neurons in both compartments is used as a weight.

The key feature of the model that relates the estimates of hierarchy to the biological measure (i.e. the transformed SLN values) is the incidence matrix, ***X***, of the cortical graph. The incidence matrix of the graph is defined to have a column for each node and a row for each link. In each row, all values are 0 except for the two nodes of the link, taking on values of −1 and 1, respectively for source and target, if the link is directed. The product of the incidence matrix and the vector of hierarchical values, *h*, maps the differences in hierarchical value between two areas with the differences between the transformed SLN, such that:

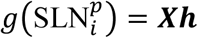

where the left side of the equation is the difference between transformed SLN values of the source area *i* and the injection site *p*. The vector *h* contains the hierarchical coefficients to estimate and its length is equal to the number of columns of the model matrix. The model as defined is not identifiable because the sum of every row equals 0, but by dropping one column, for example, area V1 in the visual hierarchy, the hierarchical coefficients can be estimated with the area corresponding to the dropped row fixed at a hierarchical level of 0.

This resembles a logistic or probit regression problem. However, these models yield standard errors for the estimated coefficients that are unrealistically small. Alternatively, we have used a beta-binomial model; this arises naturally as a binomial model in which the response, which in the current context is the expected SLN value, has variability greater than that specified by the binomial distribution and its distribution is described by a beta distribution (Lesnoff and Lancelot, 2012). For present purposes, the model can be reparameterized to include a dispersion parameter that better models the overdispersion typically observed in neural counts (see (Markov et al., 2014a) for further details). Once the statistical model is specified, the coefficients are estimated by maximum likelihood. Note that because numbers of neurons are used in the model and not just the SLN proportions, this method generates a weighted hierarchy.

The formalization is quite general. For example, if instead of SLN, a binary variable is used simply specifying whether a connection is of a FF or FB type, then the model corresponds to the original problem that Felleman and Van Essen solved. We have found that fitting the model in this fashion leads to coefficients with much larger standard errors (**Figure 7A**), thus, providing an alternate demonstration of the indeterminacy or more limited information on hierarchy contained in purely binary relations. Thus, the use of a quantitative measure of hierarchy leads to a more exact solution (**Figure 7B**).

**Figure 7.**
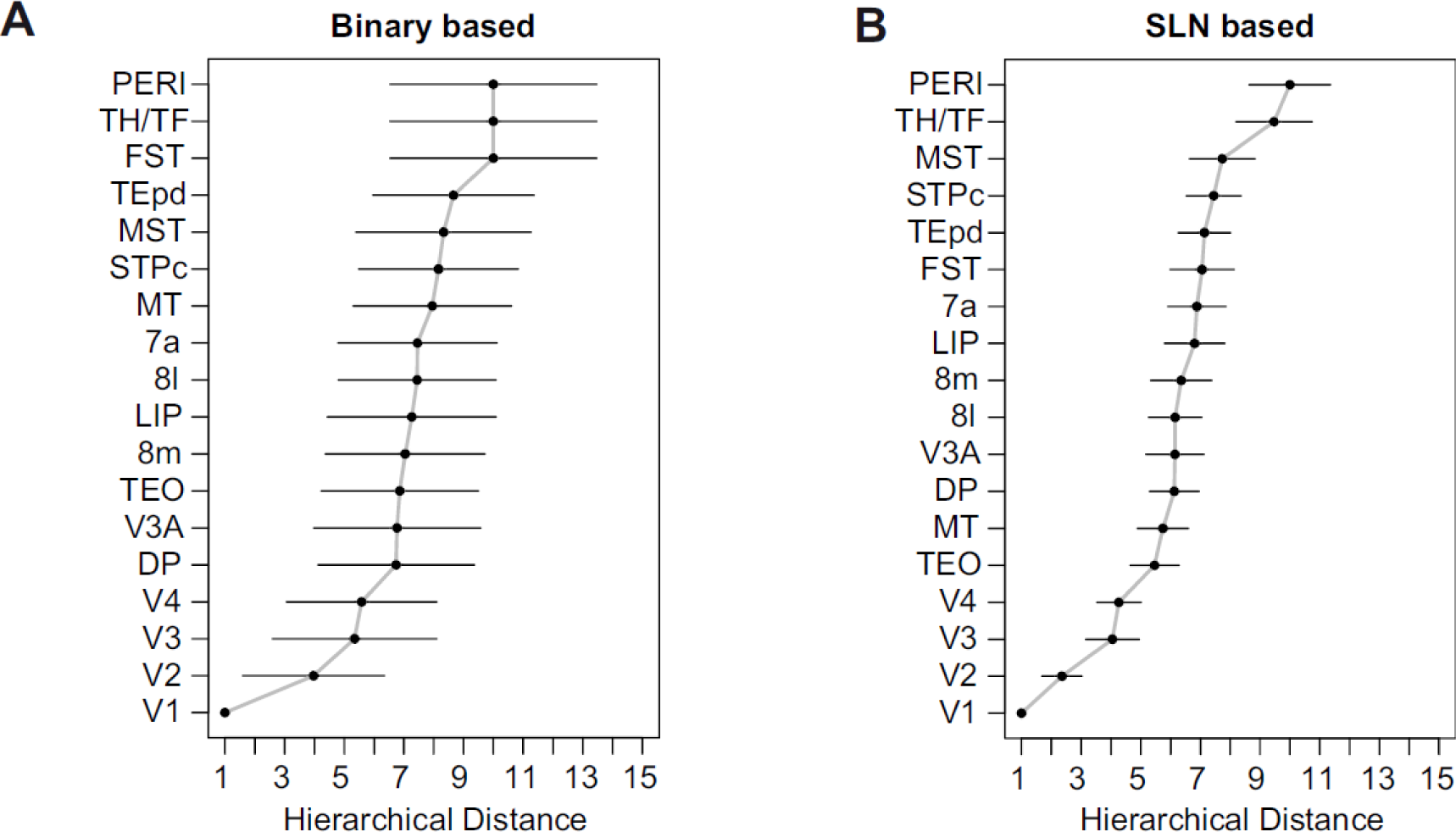
Precision of estimated hierarchy based on hierarchical index. **A**. Estimated hierarchy obtained using logit transformed SLN values as a measure of hierarchical distance. The counts of supra- and infragranular neurons are used as weights. The error bars are 95% confidence intervals estimated from the covariance matrix of the fitted model. **B**. Estimated hierarchy using a binary variable as an indicator of the hierarchical relation between area pairs. A logit link was also used in this case. The larger 95% confidence intervals demonstrate the loss of precision in estimating the hierarchical distance when using only binary information about connectivity.

To summarize, a qualitative assessment of a hierarchical gradient is initially evaluated visually by choosing a (possibly transformed) measure of the hierarchical difference between area pairs and using pairs plots to assess the pairwise consistency of the distance measure. If the evidence looks promising, the hierarchical values are obtained by fitting a model that maps the hierarchical estimates to the biological measure of the gradient via the network incidence matrix. If a suitable probability distribution can be assigned to the biological measure, the solution can be obtained by maximum likelihood, but other methods, for example introducing Bayesian priors, might be applied in appropriate circumstances.

The visual hierarchy estimated from our model is shown in **Figure 8A** and resembles qualitatively the Felleman and Van Essen model (Felleman and Van Essen, 1991). In contrast, the levels are continuously distributed. Here we have split FEF into area 8L and 8M corresponding to regions involved in small and large saccades, respectively. Area 8L occupies a low position in the hierarchy while 8M is several levels higher. The goodness of fit of the model is indicated by plotting the empirical SLN values against those predicted by the model (**Figure 8B**) and shows that the model accounts for a high proportion of the variance in the data. The functional implications of this model have been explored in several contexts (Bastos et al., 2015b; Chaudhuri et al., 2015; Magrou et al., 2018) and observations indicate that it is applicable to anterograde tracing data from the mouse (D’Souza et al., 2020).

**Figure 8.**
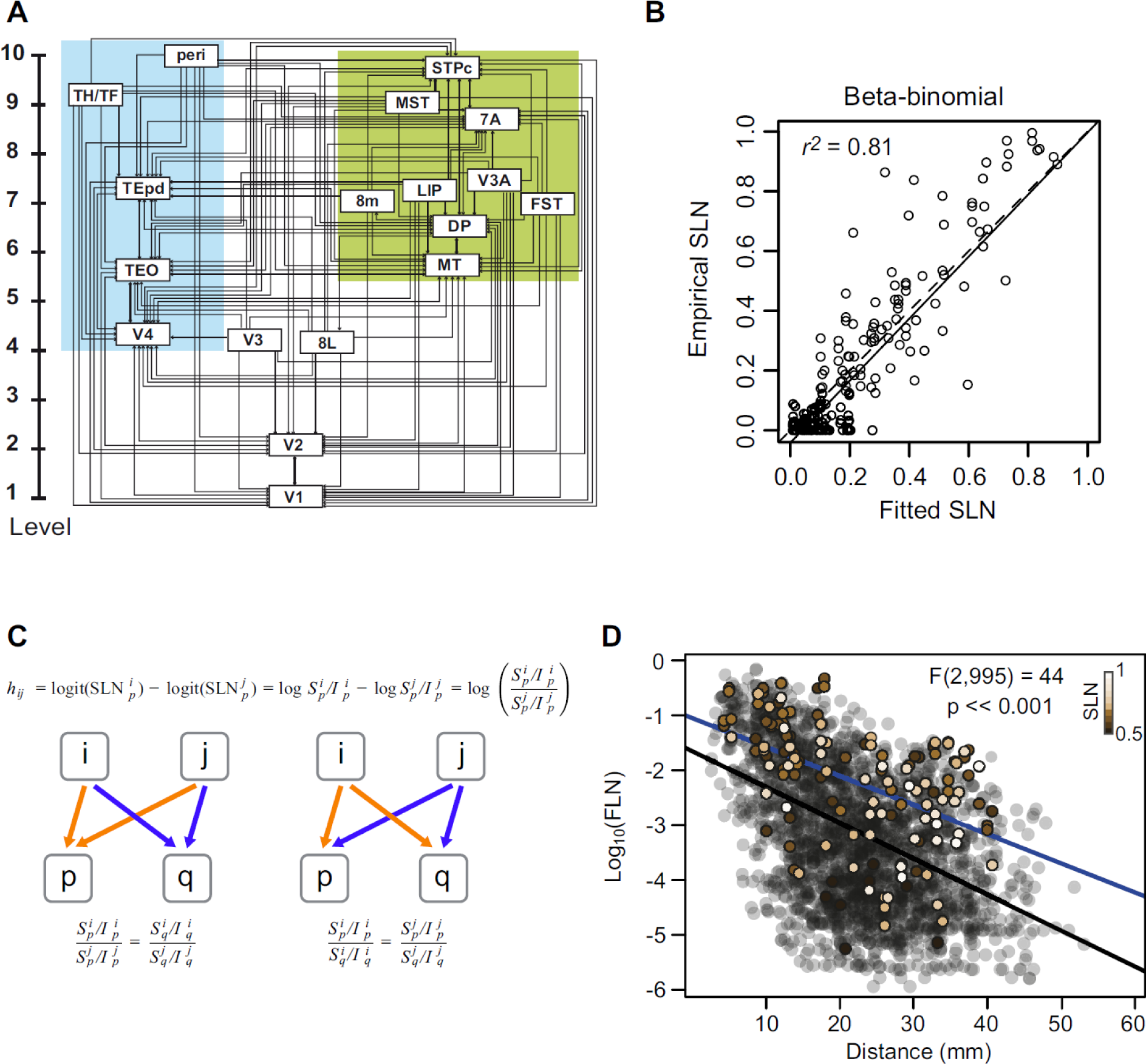
Hierarchical organization of visual areas (A) estimated from the beta-binomial model. The model only provides the vertical level of the areas with respect to the lowest level. For clarity of presentation, we have separated them laterally into ventral and dorsal stream areas. The estimated values are only unique up to adding a constant and multiplying by a coefficient. Here, we have the areas to span the range 1-10. **B**) The scatter plot shows the empirical SLN values plotted against those predicted by the model. The solid line is the unit slope line through the origin and the dashed line is the best fit linear regression. **C) Hierarchical distance**. The hierarchical distance, *h*_*ij*_, between common projections from areas *i* and *j* to area p, defined as the difference of logits of their SLN values, is equivalent to the log of the ratio of their supra- to infra-granular projection strengths to area *p*. a. This definition implies that the ratio between the laminar ratios of areas *i* and *j* to area *p* (orange arrows) is the same as that for any other target area *q* receiving projections from the same source areas (blue arrows), as formalized in the equation below the diagram. This is because the hierarchical distance from *i* to *j* should be the same for injections in both areas *p* and *q*. b. A rearrangement of the equation (below) implies, also, that the ratio between the laminar ratios of projections from a common source area, *i*, to areas *p* and *q*, will be the same for any other common source area, *j*, to the same target areas. **D) Cortical-cortical strong loops**. The strength-distance relation of 1615 projections from 91 to 29 cortical areas obtained from retrograde tracer injections. The transparent black points indicate all of the projections except those that participate in strong-loops in beige. The color gradient on these symbols corresponds to SLN strength as indicated by the inset color bar. The black line is the best fit linear regression to the transparent black points and the blue line is the best fit to the strong-loops. The F-statistic indicates the result of a nested likelihood test indicating the probability of a difference in strength between the two sets of points as large as that obtained under the null hypothesis that the true difference is zero, when physical distance via the WM is taken into account.

The use of a transformation of SLN to estimate hierarchical distance imposes strong constraints on inter-areal SLN relations. We demonstrate this in **Figure 8C** by using the logit transform, which performs quantitatively nearly identically to the probit transform (**Figure 4C**), but allows expression of the hierarchical relations in terms of ratios of projection strengths. The model defines hierarchical distance, *hij*, between two areas, *i* and *j*, as the difference between the logit of SLN values for their projections to a target area, p, shown in the top equation. For the logit, this distance is just the natural log of the ratio of ratios of supra- to infragranular projection strengths from areas *i* and *j* to p (orange arrows in **Figure 8C**, left). If the hierarchical distance measure is consistent, we expect that ratio to be the same for projections to any other area, q, (blue arrows in **Figure 8C**, left) as shown by the equation below the diagram. A simple rearrangement of this identity demonstrates that the ratio of projections from area *i* to areas p and q (orange arrows in **Figure 8C**, right) should be the same for any other area *j*, projecting to areas p and q. Thus, the hierarchical model we propose implies strong invariances in the ratio of FF to FB projection strengths from common inputs and outputs across areas. We further hypothesize that these invariances impose constraints on the exchange and stability of information flow across the cortical hierarchy.

One might suppose that when simultaneous retrograde tracer injections are made in reciprocally connected areas that the pair of areas would display a reciprocal FF-FB relation. That is to say, the origin of the majority of projections from one area would arise from upper layers and the principal origin of the reciprocating projections from lower layers. This arrangement would naturally lead to the hierarchical regularities that we observe. However, this regularity is not imposed by our model, nor is it always found to occur. In effect, this is what explains the surprising observation (noted above in the hierarchy derived in **Figure 8A**) of a prefrontal area like 8L at the same hierarchical level as early visual areas V3 and V4. As expected, the projections from several lower order visual areas, e.g., V4 and TEO, to area 8L originate in upper layers signifying FF, projections. However, 8L projects back to these areas through FF projections also originating in upper layers (Barone et al., 2000; Markov et al., 2014b). We designate such FF-FF connections as strong loops (Crick and Koch, 1998). They correspond to the situation described earlier in an order relation that when *A*≥*B* and *B*≥*A*, then *A*=*B*.

In the Felleman and Van Essen model, FF (FB) projections connect upstream (downstream) areas in a bottom-up (top-down) fashion placing hippocampus, anterior temporal and prefrontal areas at the top and occipital areas at the bottom of the visual hierarchy. As such, this model described two main counterstreams involved in hierarchical processing within the visual system: a rostro-caudal gradient of FB projections reciprocated by a caudo-rostral gradient of FF projections. Interestingly, in the data base of visual areas on which the Felleman and Van Essen model was based, only one pathway was reported – between FEF and CITd (dorsal part of inferior temporal cortex) that corresponds to a strong loop. This led Crick and Koch (Crick and Koch, 1998) to speculate that such configurations were forbidden in cortical organization. However, we have identified a significant number of strong loops in our data base. **Figure 8D** shows how the log_10_ FLN varies as a function of distance estimated through the white matter from the source area to the target injection site, referred to as white matter (WM) distance), replotted from Ercsey-Ravasz et al. (Ercsey-Ravasz et al., 2013). The beige points correspond to those connections that participate in strong-loops i.e. area pairs for which SLN is greater than 0.5 in both directions. For reference, the saturation of these points indicates their SLN values with less saturated points indicating a higher SLN, as shown by the inset color bar. As expected, most of the SLN values near 1 cluster to the right as more long distance connections. There is a fair amount of scatter among the points but they do display a distance rule (blue regression line) just as the rest of the connections do (black regression line). Interestingly, the strength of the strong loop projections is on average greater than the rest of the projections. This suggests that they are likely to play a unique role in cortical computation. What that role is currently remains a mystery. However, there is experimental evidence in favor of an attentional role concerning the strong-loop between FEF and V4. FEF projections can drive attentional response in V4 (Moore and Armstrong, 2003), through selective gating of V4 receptive-fields (Armstrong et al., 2006). Further evidence points to the involvement of fast rhythmic synchronization during FEF attentional modulation of V4 visual responses (Gregoriou et al., 2012; Gregoriou et al., 2009), strongly suggesting a supragranular origin (Bastos et al., 2015a; Markov et al., 2014b).

The results reveal a high-degree of consistency of the structural regularities underlying cortical hierarchy in the sensory cortex. But how generalizable are these findings across the more anterior regions, particularly in frontal and prefrontal cortex (Badre and D’Esposito, 2009; Choi et al., 2018)? One of the few studies that has addressed the structural hierarchy with tract tracing of the prefrontal cortex (Goulas et al., 2014) found little evidence of the rich rostral to caudal hierarchical organization that has been reported in human imaging studies (Badre and D’Esposito, 2007; Koechlin et al., 2003). The controversial issue of frontal cortex and hierarchical control shows promise of resolution via a revision of the concept of a unitary model ensuring a unidimensional gradient. Recent reports favor distinct networks that are proposed to interact thereby ensuring a global hierarchical structure (Schumacher et al., 2019). Nevertheless, the mismatch between the multistage cascade architecture mediating a temporal organization of cognitive control and inter-areal connectivity contrasts with the situation in the visual cortex where there is a smooth increase in receptive field size ascending the Felleman and Van Essen cortical hierarchy (Roelfsema and de Lange, 2016). The mismatch reported in the prefrontal cortex is between the concept of a smooth and gradual rostral-caudal organization found in the imaging studies and the connectivity as found in the collated and binary data base. What about the relation of SLN across prefrontal areas? In **Figure 9A**, the SLN pairs plots for the prefrontal cortex show an encouraging agreement with that described in visual cortex. The hierarchical scale values estimated from the statistical model described above (**Figure 9B**) seem to support a rostral-caudal organization with F1 at the bottom and areas 10 and 24c (the most anterior limbic region) at the top. Note, analysis based on more complete coverage of the frontal cortex might give significantly improved results.

**Figure 9.**
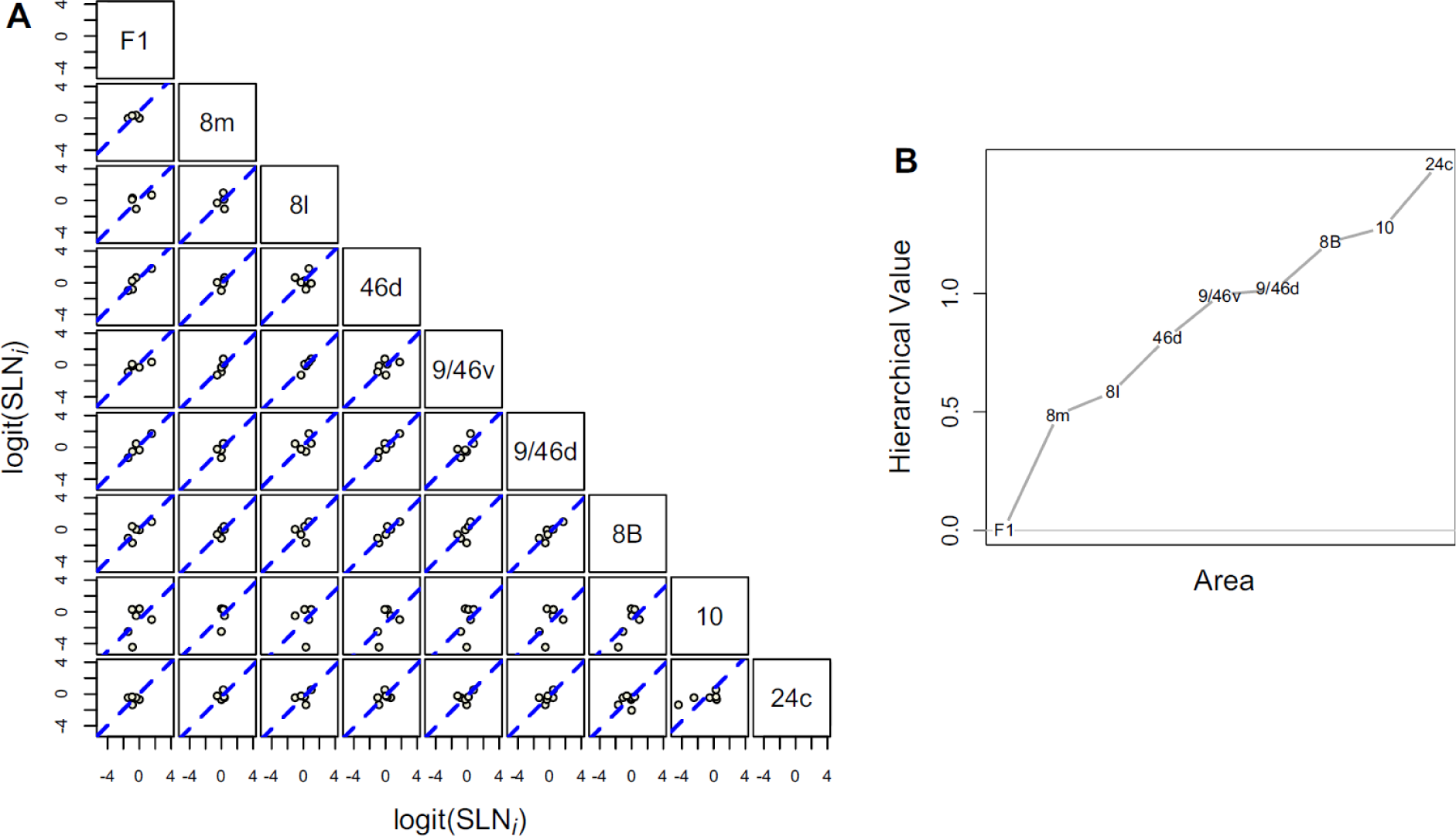
**A)** Scatter plots of logit transformed SLN values of common source areas from pairs of 9 frontal and pre-frontal areas, obtained from retrograde tracer injections. The plots follow the same format as previous in Figures 6 and 7. The dashed blue line in each plot is the best fit line of unit slope. **B)** Hierarchical scale values estimated for the 9 areas based on the proposed statistical model. Area F1 was assigned a value of 0 for model identifiability.

## 5. Dual stream Architecture and its functional significance

In the preceding section we showed that the contributions of supra- and infragranular layers in the projections across hierarchical levels were highly consistent. The consistency of our measure of hierarchical distance, in conjunction with the known differences in oscillatory properties of laminar compartments of the cortex, suggests that FF and FB interactions are not only anatomically distinct, but (i) use specific frequencies for communication and (ii) play specialized roles in cortical computation. Here we address how these functional properties relate to the detailed anatomical properties of FF and FB pathways. In the light of recent findings showing FF and FB constitute distinct pathways in both the upper and lower layers constituting a Dual Stream Architecture (Markov et al., 2014b), leads to the hypothesis that FB signals in upper and lower layers have distinct roles in information processing.

There are a number of reasons for expecting that supra- and infragranular layers might house different FF and FB pathways. During corticogenesis the supragranular compartment is generated by a primate-specific germinal zone (Smart et al., 2002), exhibiting uniquely complex lineages (Betizeau et al., 2013; Dehay et al., 2015; Lukaszewicz et al., 2005), findings that have consequently been amply confirmed in human corticogenesis (Geschwind and Rakic, 2013). These specialized developmental origins of the supragranular layers are linked to the observed expansion of these layers in primates culminating in human (Cahalane et al., 2014; Sousa et al., 2017), and a diversification of cell-types, which we speculate underlies the observed coding properties of these layers in the adult primate cortex (Tang et al., 2018; Vinje and Gallant, 2000; Wang and Kennedy, 2016; Willmore et al., 2011). A number of studies have shown that supragranular layers exhibit sparse coding in which large numbers of cells are characterized by low levels of activity and a sensory stimulus activates only few neurons (Barth and Poulet, 2012; Crochet et al., 2011; Haider et al., 2013; Harris and Mrsic-Flogel, 2013; Petersen and Crochet, 2013; Tang et al., 2018). In a sparse code information is encoded at any instant by the spiking of a small number of neurons, as opposed to a dense code where overall activity is high and information is encoded by variation in firing rate as observed in the infragranular layers (Sakata and Harris, 2009). A sparse code reduces redundancy and is arguably more efficient. Studies indicating sparse coding in supragranular layers find evidence of higher levels of recruitment of inhibitory drive in these layers via fast spiking PV+ neurons (Hu et al., 2014), which supports the presence of distinct frequency channels for FB and FF communication (Bastos et al., 2018; Bastos et al., 2015b; Michalareas et al., 2016). In addition, sparse coding, supragranular neurons in area V1 showed more complex and specific selectivity than expected for primary visual cortex (Bonnefond et al., 2017).

A more detailed description of the laminar organization of inter-areal connectivity suggests that variation of SLN with distance has complex origins concerning inter-areal connectivity in sublayers of the cortex. Exhaustive examination of inter-areal connectivity shows, that whereas canonical FB streams have been traditionally allocated to infragranular layers, a robust supragranular FB stream is in addition found in layer 2 in all areas examined in the extrastriate visual cortex of the macaque (**Figure 10A**) (Markov et al., 2014b). In addition to the layer 2 FB, we found some evidence of a layer 5 FF stream. Hence, in both upper *and* lower compartments there is a counter stream leading to the term dual counterstream architecture. Interestingly, the two FB streams in the supra and infragranular layers will impact differently the canonical microcircuit (**Figure 10B**) (Douglas et al., 1989). The strict segregation of FF and FB streams was hypothesized by Ullman in his counterstream model, which he proposed allows a bi-directional search for optimal matches between descending and ascending pathways (Ullman, 1995, 2000).

Closer examination of the individual streams showed that each obeys a unique distance rule. In all streams labeled cell counts decline with WM distance, however the rate of decline is characteristic for a given stream. In this way, the classical FB stream in layer 6 of the infragranular layers has the most gradual decline so that these FB connections span the biggest distance in the cortex. This contrasts with the layer 2 FB, which shows a much shorter space constant. Hence it is the combination of the space constants of the layers 2 and 6 FB streams that leads to the observed SLN values going up stream from near to far-distant areas (See **Figure 11**). The classical FF stream in L3 is also long-distance stream, but significantly less than the FB layer 6 stream, thus leading to the observation of the greater reach of FB pathways compared to FF pathways (Markov et al., 2014b).

**Figure 10.**
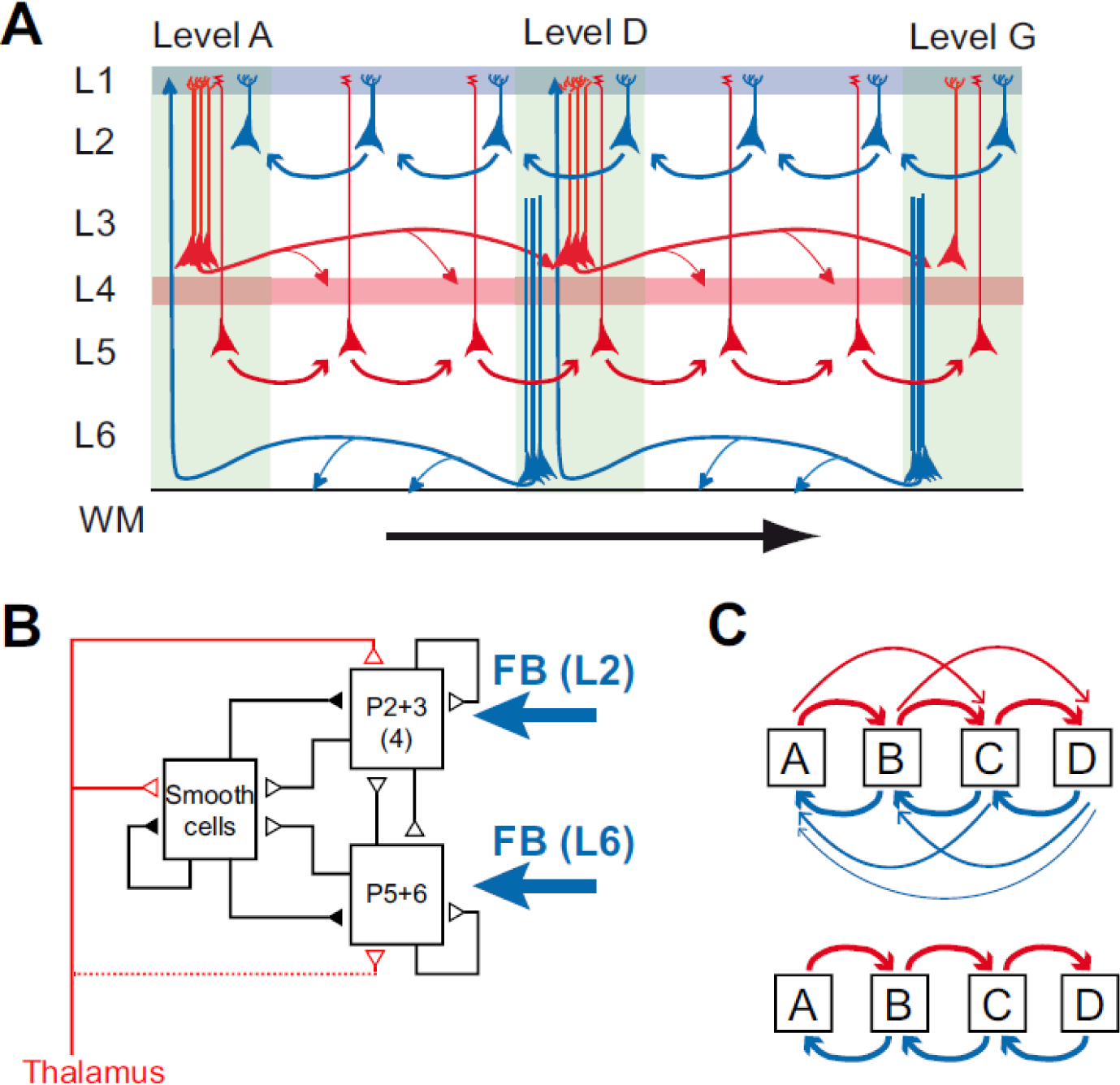
Dual counterstream architecture of inter-areal pathways. **A)** Parent neurons at L3 and L5 have FF projections (red) to higher order areas reciprocated by FB projections (blue) in L2 and the L6. Simultaneous tracer injections in high and low areas show that the upper layer counterstream has near 100% segregation, i.e. the FF (FB) neurons do not send axon collaterals to lower (higher) order areas. However the evidence that the FF and FB pathways form continuous streams, as depicted here is indirect; what crucially remains to be elucidated are the laminar details on the connectivity and the cellular targets. **B**) the canonical microcircuit showing the two FB pathways targeting L2 and L6. Modified from (Douglas and Martin, 1991); **C)** the incorrectly assumed serial processing (lower) between areas that is not observed in the cortex, where instead each areas project to all upper and lower stream areas (all to all). (panel A from (Markov et al., 2014b));

**Figure 11.**
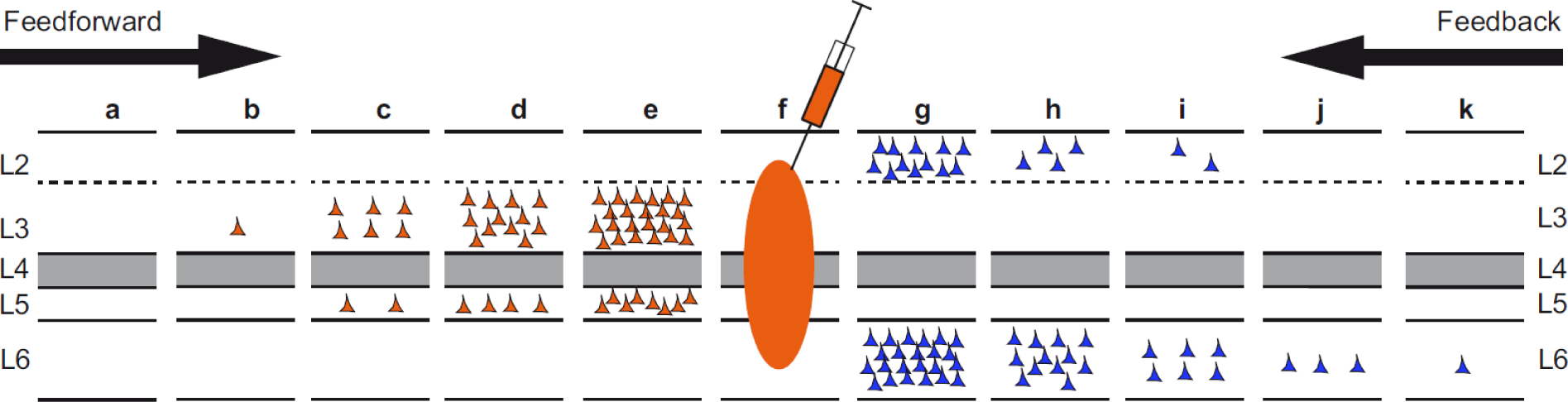
Distance effects of labeling in individual layers. This figure how FB projecting neurons are differentially distributed in L2 and 6 and FF in L3 and 5. The characteristic SLN gradient found in up- and down stream areas shown in figure 2 is due to different distance rules operating in individual layers. Hence the short-distance spread of labeled neurons in L2 coupled with the long-distance spread in L6 leads to the observed decrease in SLN with increasing FB hierarchical distances. Likewise the long-distance spread of labeled neurons in L3 coupled with the short-distance spread in L5 leads to the observed increase in SLN with increasing FF hierarchical distances.

Hence, the dual counterstream architecture shows that the relative rate of decline in individual streams determines the way that SLN is modulated by WM distance. In the previous section (Hierarchy –input consistency) we showed that the agreement between SLN values across hierarchical levels is relatively constant across the extrastriate macaque cortex, but less so for the prefrontal cortex. These differences between frontal and posterior regions could be driven by two sets of factors. Firstly, quite simply the space constant can change in individual layers so that the two regions sample supra- and infragranular layers over different WM distances. For example, if the difference in space constants of layer 2 and 6 FB streams are reduced then so are the SLN differences and there will be a reduction of the hierarchical distance as such between a group of areas with a common target. A second factor could be the identity of cells in the two streams. Comparison of the gene expression across species has revealed that some genes in rodents that are exclusively expressed in deep layers are expressed in the supragranular layers of primates (Zeng et al., 2012). Such changes in the laminar phenotype could perhaps occur across regions meaning that the layer 2 FB pathway in the prefrontal cortex may not correspond to the same identities as the FB pathway in extrastriate cortex.

## 6. Functional characteristics of FF and FB pathways

In the present review we propose that cortical hierarchy can be gauged from the *nature* of the interactions between different brain areas, in the same manner that hierarchies in social networks reflect the nature of interactions between people. Crucially, our measure of hierarchical distance shows that SLN values of area pairs are highly consistent across multiple hierarchical levels. This consistency in conjunction with the known differences in oscillatory properties of laminar compartments of the cortex suggests that FF and FB interactions are not only anatomically distinct, but (i) use specific frequencies for communication and (ii) play specialized roles in cortical computation. Here we address how these functional properties relate to the detailed anatomical properties of FF and FB pathways.

As described above functional interactions between brain areas are distance dependent (Bastos et al., 2015b; D’Souza et al., 2016; Michalareas et al., 2016; Richter et al., 2018; van Kerkoerle et al., 2014). Granger-causality was used to quantify functional FF and FB interactions, thereby allowing the strength of these interactions to be quantified for individual frequency bands. Neuronal populations show spontaneous fluctuations over time that are driven by brain rhythms in different frequency bands, such as theta (3-8Hz), alpha (8-12Hz), beta (12-30Hz) and gamma (30-80Hz). As already noted above, neocortical rhythms do not comprise band-limited sinusoidal oscillation but typically describe noisy oscillations with energy in a wide frequency range (Burns et al., 2011; Spyropoulos et al., 2020). FF Granger-causality is particularly strong in the gamma-frequency band, while FB Granger is strong in the alpha and beta-frequency band (Bastos et al., 2015b; Michalareas et al., 2016; Richter et al., 2018; van Kerkoerle et al., 2014).

The finding that Granger-causality in the FF (FB) direction is strong in gamma (alpha/beta) frequencies is partially dictated by the cellular targets of inter-areal pathways. FF inputs target both excitatory and GABAergic interneurons (**Figure 12**). Importantly, the FF projections to GABAergic interneurons target almost uniquely parvalbumin – PV interneurons, which are associated with gamma-frequency rhythms and respond to excitatory inputs with high temporal fidelity (Buzsaki and Wang, 2012; Cardin et al., 2009; Jouhanneau et al., 2018; Wang, 2010), and exhibit supra-threshold resonance at gamma-frequencies (Hasenstaub et al., 2005; Pike et al., 2000). Moreover FF projections skip the infragranular layers that are canonically associated with alpha and beta rhythms (Bollimunta et al., 2008; Buffalo et al., 2011; van Kerkoerle et al., 2014). By contrast, FB projections target multiple classes of GABAergic interneurons. Of these, somatostatin – SSt interneurons are associated with the generation of slower brain rhythms (Moore et al., 2010), and can directly influence activity of neurons in infragranular compartments, which are associated with alpha/beta rhythms.

**Figure 12.**
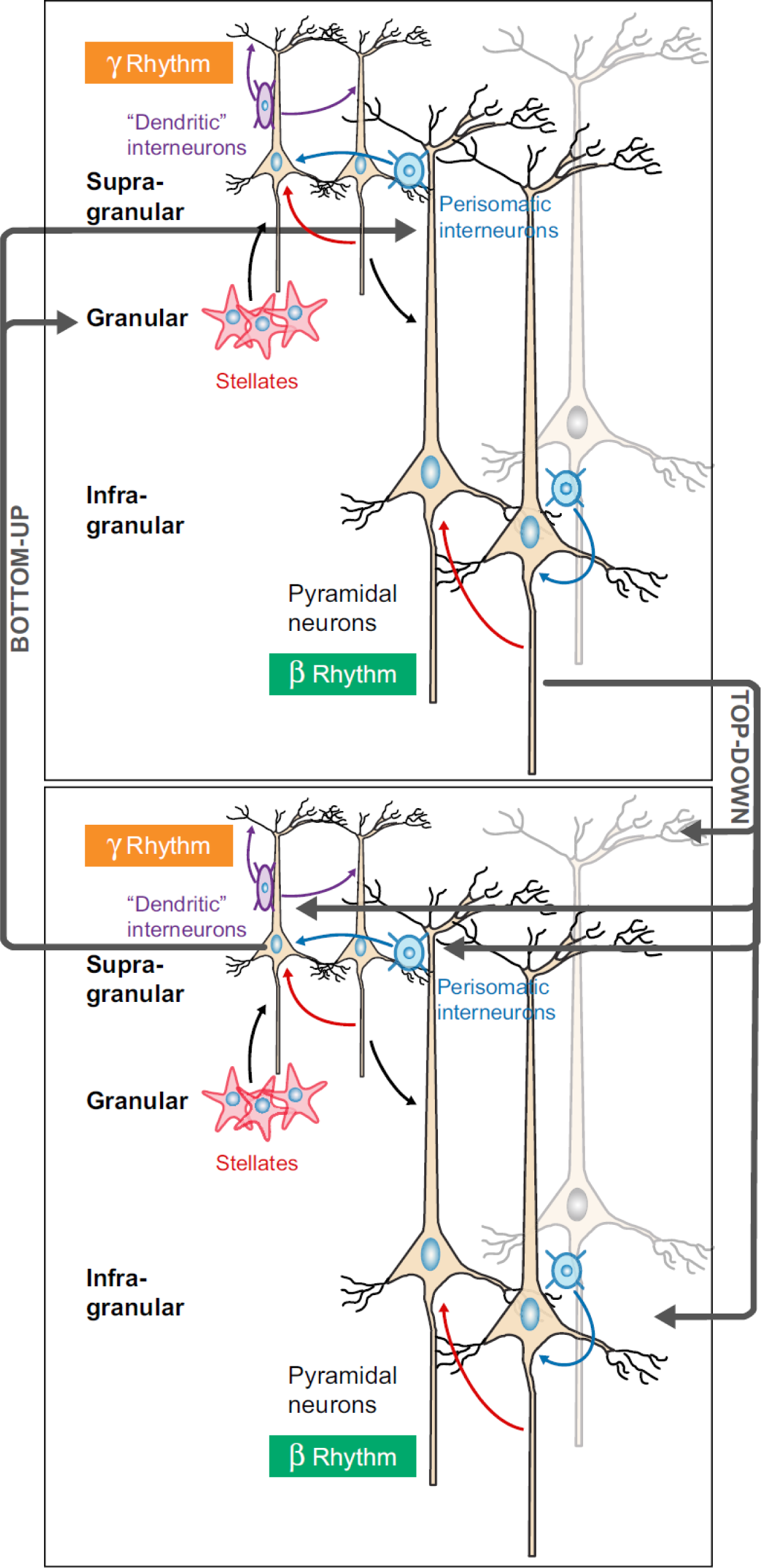
Schematic circuit for the interplay between bottom-up and top-down signaling characterized by differential frequency-band synchrony. In a reciprocally connected loop between a sensory-type area and a cognitive-type area, neural circuits in the superficial layers are endowed with strong intrinsic synaptic connections and generate stochastic oscillations in the gamma frequency range, whereas the deep layers have a propensity to display slower oscillations in the lower beta or alpha frequency range. Top-down projections originate in the deep layers and innervate pyramidal cells (brown), as well as dendrite-targeting (purple) and perisoma-targeting (blue) inhibitory interneurons. In this scenario, low beta/alpha oscillations are directly involved in top-down signaling, which interacts with locally generated gamma oscillations. Adopted with permission from Wang (Wang, 2010).

Frequency-specific information transmission can also be predicted by the properties of the *sending* and *receiving* areas, so that the global distribution of brain rhythms across areas and layers contribute to the nature of functional interactions in the FF and FB directions. Gamma-rhythms are particularly strong in early visual areas *and* superficial layers (Bastos et al., 2015a; Buffalo et al., 2011; van Kerkoerle et al., 2014; Vinck and Bosman, 2016), whereas beta rhythms are most prominent in fronto-parietal areas *and* infragranular compartments (Bastos et al., 2015a; Buffalo et al., 2011). Consequently, one expects Granger-causal FF influences from early visual areas to parietal areas to be dominated by gamma frequencies, whereas FB influences to visual areas to be dominated by beta frequencies, in agreement with the fact that major long-range FF projections originate nearly exclusively from superficial layers, and FB from infragranular layers. Further, we note that gamma frequencies are generally associated with cortical activation, e.g. a visual stimulus or optogenetic activation, whereas alpha and beta frequencies are not clearly associated with a FF stimulus drive, consistent with the suggestion that FF provide the major driving influence (Covic and Sherman, 2011; De Pasquale and Sherman, 2011).

Above we hypothesized that these frequency specific channels constitute functional analogues of the SLN projections that we have quantified, leading to the expectation that they exhibit similar hierarchical properties. Bastos et al. (2015) defined the multiple Directed-influence Asymmetry Index (mDAI) based on the difference of gamma and theta vs beta frequency Granger-causality, obtained from ECog recordings in macaque. mDAI enabled these authors to reconstruct hierarchies that closely resembled the structural hierarchy derived from SLN values. The feasibility of accounting for their data with the model that we presented above is demonstrated by the strong tendency of mDAI values corresponding to common projections to lie along lines of unit slope (**Figure 13**). The mDAI index is already on a scale that appears to show the consistency necessary to apply the model, so the hierarchical values could be estimated from the incidence matrix in a global fashion, as described above, rather than by the averaging method used by Bastos et al. (2015).

**Figure 13.**
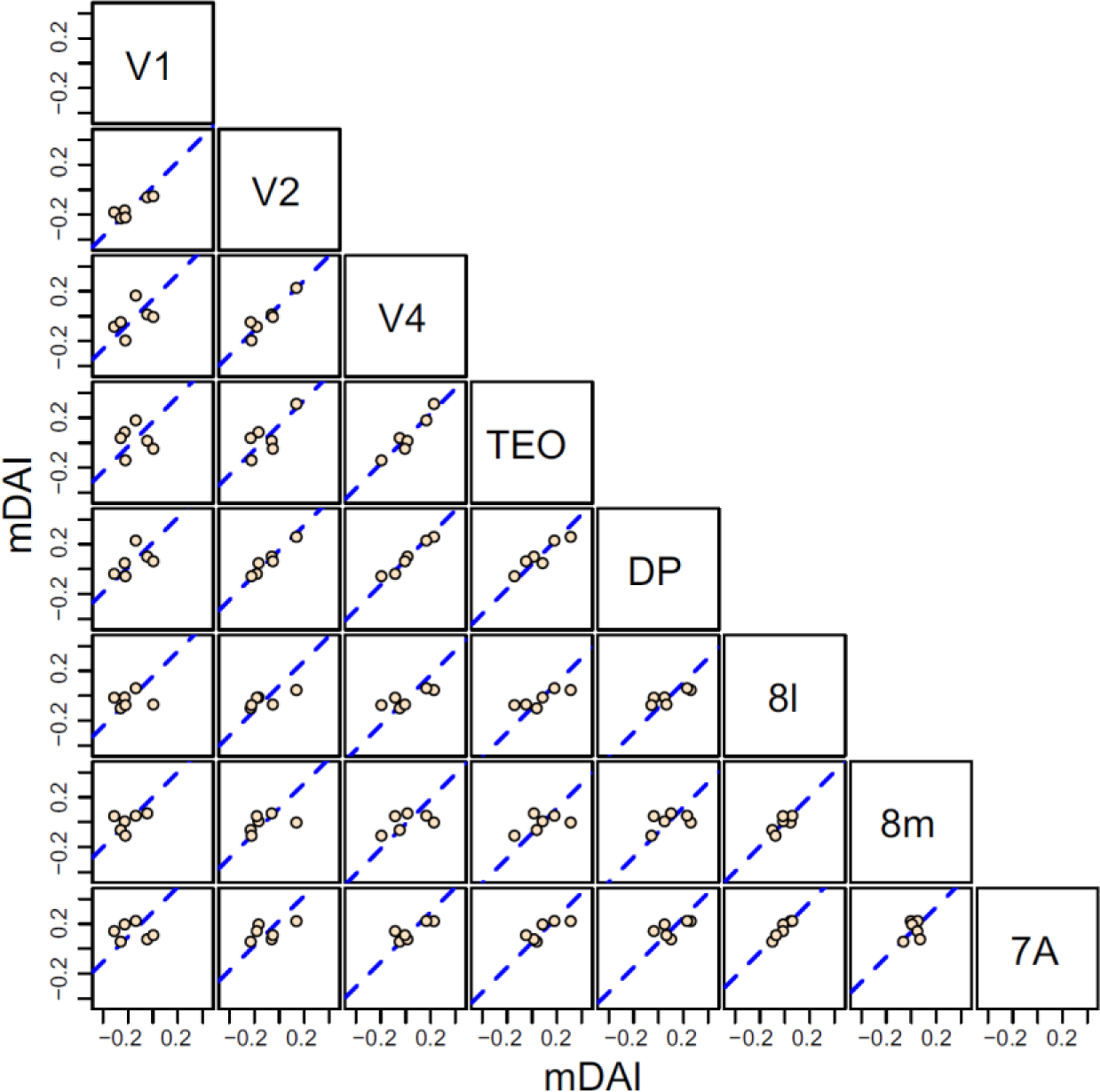
Scatter plots of a hierarchical measure of cortical distance (mDAI) derived by Bastos et al., 2015 of common source areas for pairs of 8 visual areas obtained from contrasting Granger Causality measures in gamma, theta and beta bands. The abscissa of each graph corresponds to the value calculated for the area at the top of the column and the ordinate to the area at the right of the row. Dashed blue line in each plot is the best fit line of unit slope.

Future work will need to specifically address the nature of layer-specific functional interactions for individual FF and FB projections. The dual counterstream architecture postulates that a prominent short-range FB projection originates from L2 in the higher area. Consequently, we predict that FB from L2 is particularly strong at gamma-frequencies. Likewise, L5 is postulated to have a short-range FF projection, and to exhibit strong oscillations at alpha/beta frequencies. Hence, based on the dual counterstream architecture we predict a greater diversity of functional interactions than suggested by previous work, the elucidation of which will require multi-layer high-density recording across multiple cortical areas.

## 7. The predictive brain and the importance of topdown generative networks

A large body of physiological studies has shown that FF pathways ascending the hierarchy generate increasingly complex representations of the world in higher areas, leading to the large range of receptive field properties observed at different levels of the hierarchy. Thus, at its core, convergent FF projections carry information from the outside world, and allow this information to be gradually transformed to low-dimensional representations that can be exploited for behavior. In this respect, it is worthwhile noting that the recent success of deep neural network architectures in solving complex tasks similarly demonstrates the power of FF transformations in computation (LeCun et al., 2015; Richards et al., 2019) e.g. by forming increasingly complex representations along the feedforward hierarchy in convolutional networks (Yamins and DiCarlo, 2016).

In contrast to FF-pathways, the neurobiology of the twice as numerous FB pathways (Markov et al., 2014a) remains elusive, forming a major impediment to understanding the brain. A clearly defined role of FB connections is proposed for attentional modulation, but FB pathways are likely critical in a host of complex operations including: the comparison of internally generated predictions of sensory input with actual inputs; imagining sensory-like representations from concepts of e.g. visual objects; carrying out mental simulations and finally gating synaptic plasticity. An early conceptualization of hierarchical processing in the cortex conceived of FF pathways as driving target areas, whereas FB pathways would merely exert a modulatory influence (Klink et al., 2017); some researchers, however, proposed a more nuanced view (Bastos et al., 2012). Indeed, the simple dichotomy of the roles of FF and FB pathways is difficult to reconcile with the multiple physiological effects that are imputed to FB control. For example, in the case of perceptual completion (e.g. in illusory figures) or visual imagination, FB is usually conceived to *enhance* neural activity; by contrast in the case of filtering out self-generated sensory inputs, FB activity is expected to *suppress* neural activity. These forms of enhancement and suppression represent essentially distinct computational operations: a central concept is that of *gain modulation*, where an existing sensory representation is preserved, but in essence multiplied or divided, as in attentional mechanisms (McAdams and Maunsell, 1999). However, in the case of imagery, one expects FB to *“write-in”* a pattern of neural activity, i.e. operate a *driving* process, or alternatively selectively modulate the activity of specific spontaneously active cell assemblies. In the case of *cancelling out* self-generated sensory inputs through FB (as in self-tickling), FB activity is thought to be *subtractive* (Bastos et al., 2012). Finally, FB activity has been conceived to mediate *error* signals, playing a key role in shaping perceptual learning and synaptic plasticity. The notion of FB as a “swiss-army-knife” contrasts with FF processing which is essentially homogeneous and driving, as captured by the architecture of deep neural networks. These considerations underline the diversity of FB processes, which could be mediated by distinct neural circuits. In particular, we hypothesize that laminar and distance-determined diversity of FB pathways will exhibit anatomo-functional properties that characterize the cortical circuits underlying the diverse global influences engaged in different cognitive functions. Given the diversity of FB pathways, and the many functions in which FB is implicated, it is a daunting task to develop a unified theory of the function of cortical FB. Yet, our understanding of the brain depends crucially on the development of precise theories of cortical FB.

The core feature of FB that distinguishes them from FF is that their projections are more divergent; i.e. they project from few to many neurons. Interestingly, divergent projections are a core feature of the most popular kind of artificial neural network architectures, which are also rapidly becoming a mainstream model of sensory processing in the brain (LeCun et al., 2015; Richards et al., 2019). In FF (deep) neural networks, divergent error signals lead to an adjustment of synaptic weights of FF projections (“backprop”). In other words, in FF (deep) neural networks, the exclusive role of FB is to improve the data transformations implemented by the FF projections. For a biological organism, error signals could be provided for instance by multi-sensory signals or reward signals. However, it is an open question as to how FB would be able to adjust synaptic weights of FF projections (Whittington and Bogacz, 2019). A candidate pathway is the multiple FB projections converging onto layer 1 (Cauller, 1995). Here FB projections impact the distal dendrites of pyramidal neurons, activating non-linear NMDA-R-dependent dendritic integration mechanisms and voltage-gated calcium channels. A recent study provides evidence that top-down FB to layer 1 might indeed be involved in perceptual learning in a primary sensory area: With reward reinforcement, mice rapidly learn a behavioral task in which their response is contingent on electric-current stimulation of S1. However, when layer 1 projections from perirhinal cortex to S1 are inhibited, mice fail to learn the same task, suggesting that top-down FB is instructive for learning (Doron et al., 2019). Another candidate pathway to modulate plasticity is the FB projection to VIP+ and SOM+ interneurons, given that SOM+ neurons can gate plasticity on the dendrites of pyramidal neurons (Batista-Brito et al., 2018).

Until very recently the dominant theory of brain function was that sensory information progressing up the cortical hierarchy undergoes successive abstractions generating increasingly complex receptive fields. In this feedforward processing model of cortex function, the characterization of the receptive fields at different levels of the hierarchy has been a neuroscience success story over the past 60 years (Hubel and Wiesel, 1962). The bottom up sensory processing driving receptive field elaboration leads to the notion of the importance of single neurons coding for perceptually significant features. According to this feature detection view the cortical hierarchy will house neurons ranging from edge detectors to the proverbial grandmother neurons (Martin, 1994). However, in recent years there has been a paradigm shift in cognitive neuroscience that takes account of the inherent uncertainty of the nature of the sensory input to the brain. In this view which goes back to Helmholtz and was later championed by Richard Gregory, making sense of input from the sensorium requires knowledge of the world, which allows the brain to develop hypotheses of the world that are tested against sensory evidence (Friston, 2010; Gregory, 1997). These hypotheses can be formalized as Bayesian inferences on the causes of our sensation and how these inferences induce perception and behavior (Dayan et al., 1995; Lee and Mumford, 2003). The implementation of inference in the cortex has more recently been recast in terms of the circuits underlying the hierarchical processing that formulate generative statistical models as predictive processing (Clark, 2013; de Lange et al., 2018; Friston, 2010; Keller and Mrsic-Flogel, 2018; Rao and Ballard, 1999).

According to the predictive processing model the brain possesses a model of the world that it seeks to optimize using sensory inputs. Predictive processing postulates that the prior information that resides at the different levels of the cortical hierarchy generate descending predictions that cascade down the cortical hierarchy, allowing interaction with bottom up information ascending the hierarchy. At each level of the hierarchy the interaction between the top-down predictions and the ascending input from the sensorium is such that only the residual, unexplained sensory information (or prediction error) proceeds to the next level. The overall outcome of the hierarchical process is prediction error minimization. Predictive processing constitutes an inversion of the classical feedforward model by proposing that descending feedback pathways provide representations of the external world, which modifies the ascending signal that now indicates aspects of the world that are unexpected. Given its radical nature impacting every aspect of cortical function, predictive processing has attracted considerable attention from experimentalists which have gone some way to providing empirical support (Bastos et al., 2012; Clark, 2013; de Lange et al., 2018; Keller and Mrsic-Flogel, 2018; Walsh et al., 2020).

Much of the complexity of predictive coding concerns the circuitry underlying the interaction of FF and FB streams and this area of research is still hotly debated (Keller and Mrsic-Flogel, 2018). However, little research has focused on testing how FB connections in the brain could serve as the substrate of a top-down generative network (Bastos et al., 2012; Hinton, 2007; Kosslyn, 1994; Mumford, 1992; Senden et al., 2019). In FF visual processing, high-dimensional sensory representations such as an image are sequentially transformed into low-dimensional representations such as object categories, represented at higher hierarchical levels. Conceivably FB pathways *invert* this process by generating high-dimensional representations starting from low-dimensional variables (Hinton, 2007). We refer to such pathways as top-down generative networks. Whereas the FF projections in convolutional networks create a convergence of information from many to few, in generative networks information is relayed by divergent projections from few to many. For instance, the perception of a red apple depends on a transformation of the image of a red apple (represented by many neurons) into high-level representations in the temporal lobe (represented by few neurons). However, if we imagine a red apple, processing should start from high-level representations (few neurons) and generate a low level neural representation of the image of a red apple (represented by many neurons).

Top-down generative networks may play distinct functional roles in several cognitive processes such as predictive processing, mental simulation, imagery or selective attention. An attractive aspect of the generative top-down network hypothesis is that all of these functions are subserved by a relatively small number of anatomical FB pathways implementing a function-specific generative network. This network would then interact with distinct cellular components in individual target areas, thereby differentially impacting ongoing ascending FF activity in distinct pathways. As an example, we can take the processes of *imagination, expectancy and attention*. In the case of expectancy (e.g. walking to the kitchen and expecting to see a red apple in the fruit basket, but not in the sink), generative networks may cause a reduction of neural response to an expected stimulus, whereas that to surprising stimuli would not be suppressed by top-down predictions. In the case of attention (e.g. searching for a red apple in the kitchen), generative networks may lead to an amplification of sensory activity when we find the stimulus that we were seeking. In the case of *imagination* (e.g. thinking of a red apple), generative networks would drive activity in lower areas based on the activation of neurons in higher areas in the absence of any sensory stimulation. Since high-level brain areas contain abstract representations of objects, how do the feedback pathways of the brain achieve the remarkable feat of generating concrete sensory representations in the mind’s eye during mental imagery? An apple is an abstract concept, yet we can generate concrete instantiations of apples by imagining for example, a Pink Lady or a green Granny Smith apple on a real or an imagined table in front of us. Remarkably, we are also not limited to imagining objects as we usually see them –it is equally possible for us to imagine a red apple that is as big as a football! Remarkably, we can generate sensory experiences in environments of which we have no experience, such as standing on Mars staring out through our space helmet at the red colored landscape in front of us. These examples illustrate a cornerstone of our hypothesis: that generative networks are competent to transform abstract concepts into concrete sensory-like representations in our mind’s eye.

Strong experimental support for top-down generative networks comes from findings in human imaging experiments showing that imagined objects lead to corresponding spatial temporal activation of area V1 (Emmerling et al., 2016; Naselaris et al., 2015; Senden et al., 2019; Thirion et al., 2006). The activation of the top-down pathway is coherent with predictive processing theory where FB pathways shape the FF pathway but not the inverse. These findings of top-down generative networks creating a sensory-like representation in early visual areas could be a particularity of the human brain, thereby providing the substrate for enhanced visual imagery supporting simulation and imagination. However, this appears not to be the case; a recent breakthrough publication from the Roelfsema lab revealed spiking activity in area V1 of a macaque with respect to an object held in working memory (van Kerkoerle et al., 2017), suggesting that the fMRI activation patterns reported in human experiments could correspond to spiking activity generated by visual imagery of imagined objects (Emmerling et al., 2016; Hinton, 2007; Naselaris et al., 2015; Senden et al., 2019; Thirion et al., 2006).

The dual counterstream architecture (see Section 5) suggests several features that are congruent with the hypothesis of top-down generative networks: *(1)* the convergence of cortical FF connections in the sensory stream is mirrored by the divergence of FB connections; *(2)* source populations of FF and FB pathways are completely separate, which has been recognized as a key requirement enabling distinct functionalities of generative top-down networks (Friston, 2018; Markov and Kennedy, 2013; Markov et al., 2014b; Shipp, 2016) *(3)* on average throughout the cortex there are twice as many FB pathways than FF projections to a given area. Studies in which human subjects are cued to generate a sensory percept also provide evidence for top-down generative networks in the cortex (Emmerling et al., 2016; Naselaris et al., 2015; Senden et al., 2019; Slotnick et al., 2005; Thirion et al., 2006).

Examining the dual counterstream suggests a possible division of labor between layer 2 and 6 FB projections: Layer 2 FB and layer 3 FF projections exhibit common features that distinguish them from layer 6 FB. Layer 2 FB and layer 3 FF are short distance, topographically organized and exhibit low rates of bifurcation; contrasting with layer 6 FB which are long-distance, diffuse and have high rates of bifurcation. Thus, the layer 2 FB system may mirror the layer 3 FF system and implement a generative top-down network in which high-dimensional sensory representations can be generated through sequential stages, starting from higher brain areas. The layer 6 FB pathway, on the other hand, may determine the way in which the layer 2 generative network interacts with the local microcircuit, sending contextual signals that reflect behavioral goals and reward signals. Based on this reasoning, we hypothesize that layer 6 FB has more modulatory effects that it exerts, for example, via targeting the apical layer 1 dendrites as well as GABAergic interneurons such as SOM+ and VIP+ interneurons that modulate the activity of local pyramidal neurons (Batista-Brito et al., 2018). Testing this hypothesis will require parallel anatomical and physiological investigations. Optogenetic techniques in non-human primates could allow the injection of precise spatiotemporal activation patterns into specific laminar compartments of higher brain areas, combined with physiological measurements of activity in lower brain areas.

### Box 1

#### Experimental Exploration of the Dual Counterstream Architecture

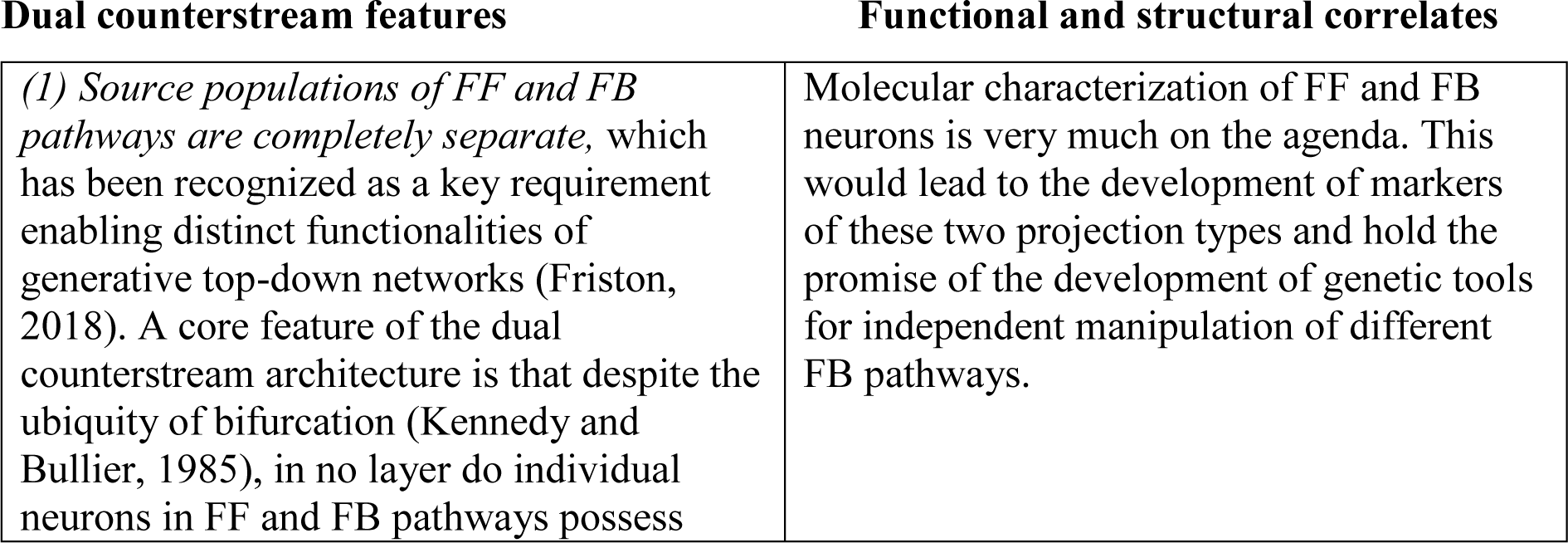
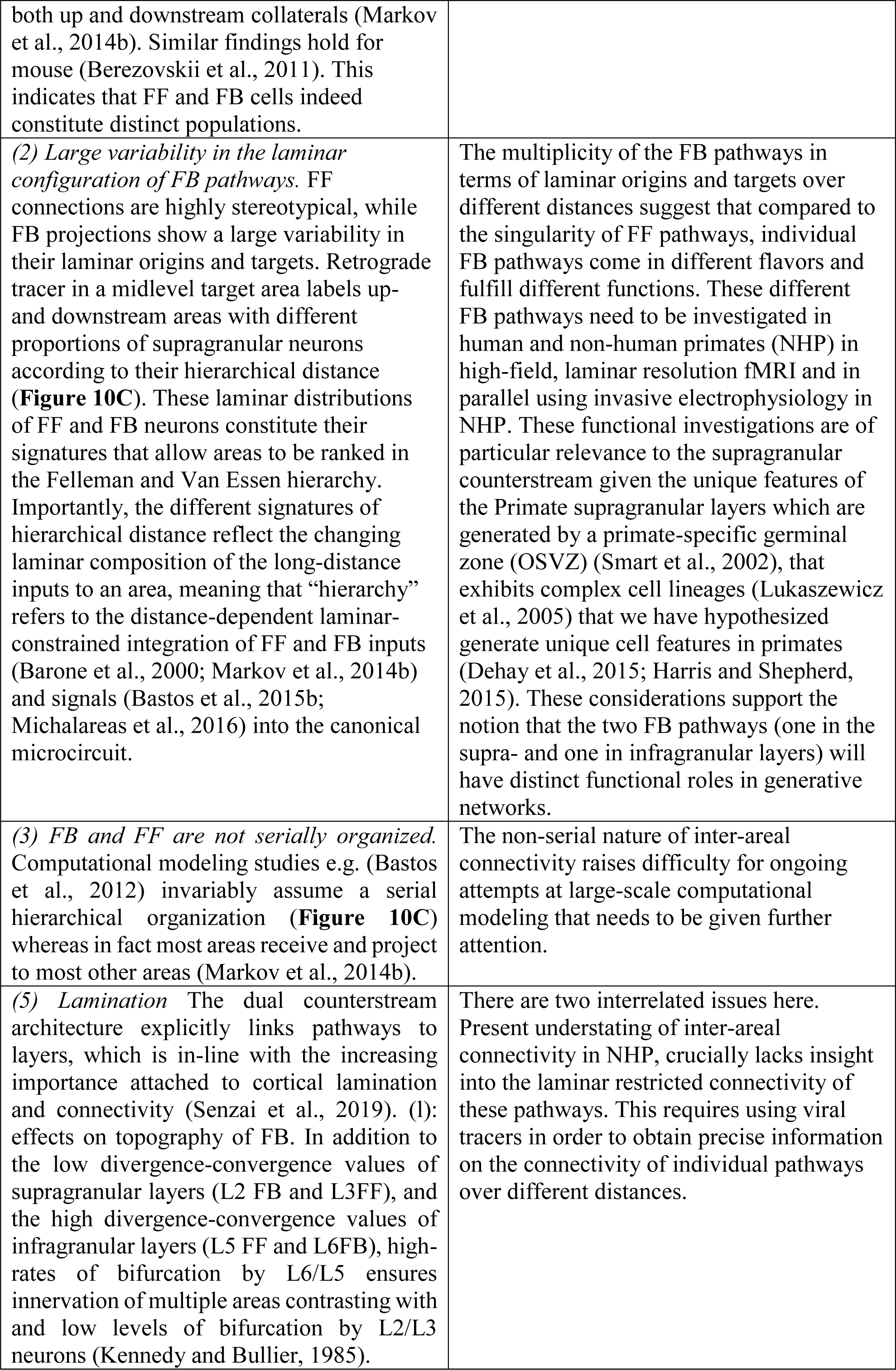
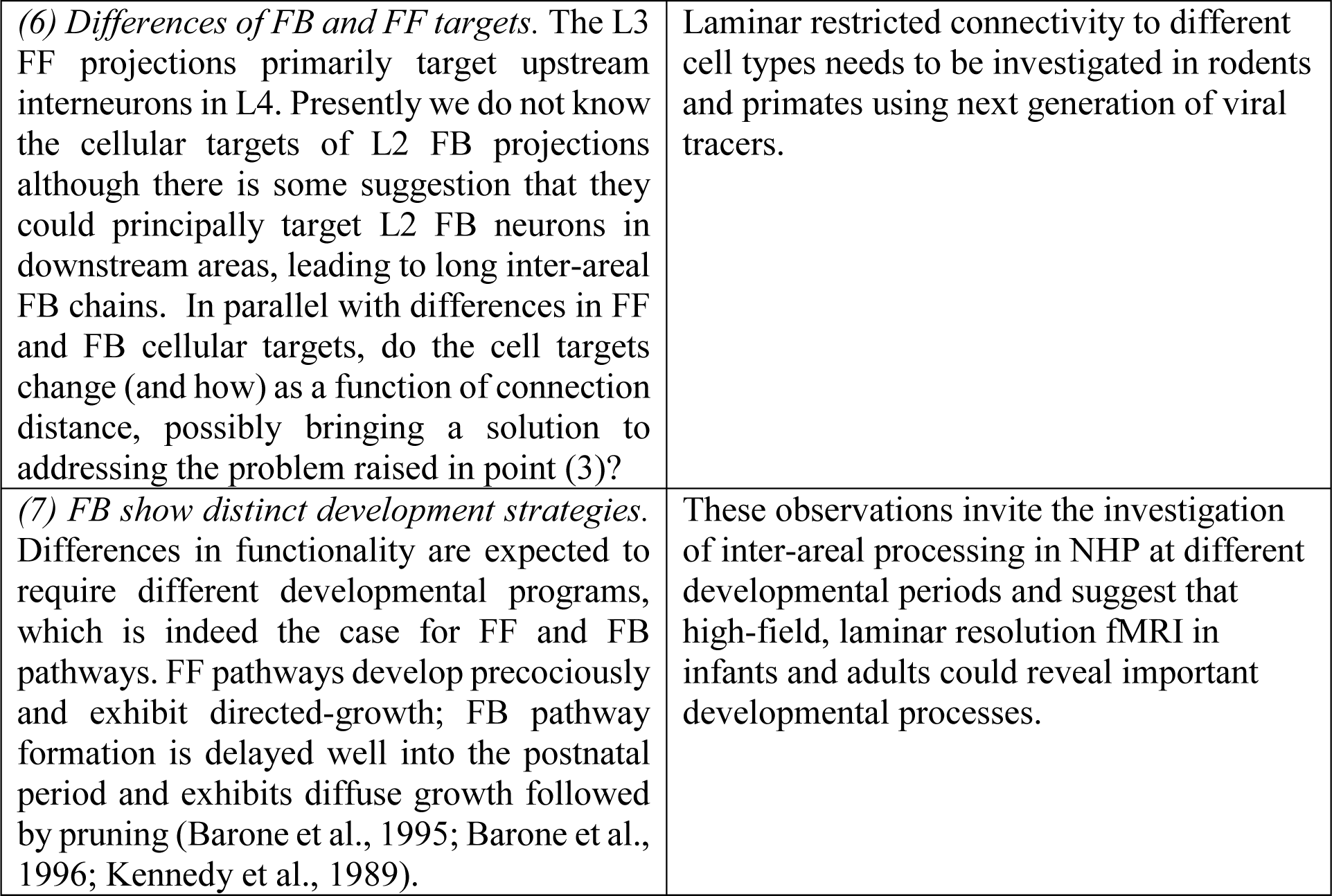

Experimental Exploration of the Dual Counterstream Architecture.
| Dual counterstream features | Functional and structural correlates |
| --- | --- |
| *(1) Source populations of FF and FB pathways are completely separate, which has been recognized as a key requirement enabling distinct functionalities of generative top-down networks (Friston, 2018). A core feature of the dual counterstream architecture is that despite the ubiquity of bifurcation (Kennedy and Bullier, 1985), in no layer do individual neurons in FF and FB pathways possess* | Molecular characterization of FF and FB neurons is very much on the agenda. This would lead to the development of markers of these two projection types and hold the promise of the development of genetic tools for independent manipulation of different FB pathways. |
| both up and downstream collaterals (Markov et al., 2014b). Similar findings hold for mouse (Berezovskii et al., 2011). This indicates that FF and FB cells indeed constitute distinct populations. |  |
| (2) *Large variability in the laminar configuration of FB pathways.* FF connections are highly stereotypical, while FB projections show a large variability in their laminar origins and targets. Retrograde tracer in a midlevel target area labels up- and downstream areas with different proportions of supragranular neurons according to their hierarchical distance (**Figure 10C**). These laminar distributions of FF and FB neurons constitute their signatures that allow areas to be ranked in the Felleman and Van Essen hierarchy. Importantly, the different signatures of hierarchical distance reflect the changing laminar composition of the long-distance inputs to an area, meaning that “hierarchy” refers to the distance-dependent laminar-constrained integration of FF and FB inputs (Barone et al., 2000; Markov et al., 2014b) and signals (Bastos et al., 2015b; Michalareas et al., 2016) into the canonical microcircuit. | The multiplicity of the FB pathways in terms of laminar origins and targets over different distances suggest that compared to the singularity of FF pathways, individual FB pathways come in different flavors and fulfill different functions. These different FB pathways need to be investigated in human and non-human primates (NHP) in high-field, laminar resolution fMRI and in parallel using invasive electrophysiology in NHP. These functional investigations are of particular relevance to the supragranular counterstream given the unique features of the Primate supragranular layers which are generated by a primate-specific germinal zone (OSVZ) (Smart et al., 2002), that exhibits complex cell lineages (Lukaszewicz et al., 2005) that we have hypothesized generate unique cell features in primates (Dehay et al., 2015; Harris and Shepherd, 2015). These considerations support the notion that the two FB pathways (one in the supra- and one in infragranular layers) will have distinct functional roles in generative networks. |
| (3) *FB and FF are not serially organized.* Computational modeling studies e.g. (Bastos et al., 2012) invariably assume a serial hierarchical organization (**Figure 10C**) whereas in fact most areas receive and project to most other areas (Markov et al., 2014b). | The non-serial nature of inter-areal connectivity raises difficulty for ongoing attempts at large-scale computational modeling that needs to be given further attention. |
| (5) *Lamination* The dual counterstream architecture explicitly links pathways to layers, which is in-line with the increasing importance attached to cortical lamination and connectivity (Senzai et al., 2019). (1): effects on topography of FB. In addition to the low divergence-convergence values of supragranular layers (L2 FB and L3FF), and the high divergence-convergence values of infragranular layers (L5 FF and L6FB), high-rates of bifurcation by L6/L5 ensures innervation of multiple areas contrasting with and low levels of bifurcation by L2/L3 neurons (Kennedy and Bullier, 1985). | There are two interrelated issues here. Present understating of inter-areal connectivity in NHP, crucially lacks insight into the laminar restricted connectivity of these pathways. This requires using viral tracers in order to obtain precise information on the connectivity of individual pathways over different distances. |
| (6) *Differences of FB and FF targets.* The L3 FF projections primarily target upstream interneurons in L4. Presently we do not know the cellular targets of L2 FB projections although there is some suggestion that they could principally target L2 FB neurons in downstream areas, leading to long inter-areal FB chains. In parallel with differences in FF and FB cellular targets, do the cell targets change (and how) as a function of connection distance, possibly bringing a solution to addressing the problem raised in point (3)? | Laminar restricted connectivity to different cell types needs to be investigated in rodents and primates using next generation of viral tracers. |
| (7) *FB show distinct development strategies.* Differences in functionality are expected to require different developmental programs, which is indeed the case for FF and FB pathways. FF pathways develop precociously and exhibit directed-growth; FB pathway formation is delayed well into the postnatal period and exhibits diffuse growth followed by pruning (Barone et al., 1995; Barone et al., 1996; Kennedy et al., 1989). | These observations invite the investigation of inter-areal processing in NHP at different developmental periods and suggest that high-field, laminar resolution fMRI in infants and adults could reveal important developmental processes. |

## 9. Conclusion and speculations

We have shown that cortical hierarchy can be defined by connectivity gradients and the functional correlates of these gradients. In other words, inputs to a cortical area sample the cortical sheet in a principled manner defined by the differential space constants of the distance rules governing the individual cortical layers. This approach to understanding hierarchy is anchored in the recognition that it is the activity and connectivity linking neurons across the cortex that will ultimately reveal the process underlying the dynamics of cortical function. Link based investigation is complemented by characterization of the nodes. Helen Barbas has championed the structural model, which shows that laminar differentiation allows hierarchical ranking of cortical areas that correlates well with connectivity patterns (Barbas, 2015). The structural model is of particular interest because it allows ranking of cortical areas via gradual variations of cytoarchitecture and myeloarchitecture (Sanides, 1972), and has opened the interpretation of large-scale models of the cortex to investigation with non-invasive imaging techniques that can be applied to the human brain (Burt et al., 2018; Margulies et al., 2016; Paquola et al., 2019).

The central argument of this review is that cortical hierarchy can be usefully thought of as the gradual changes in the cortical input requirements of the local cortical circuit that in terms of synaptic mass constitutes the powerhouse of the cortex. Understanding the cellular mechanisms underlying hierarchical processing require investigations of hierarchy in terms of the laminar restricted connectivity and physiology that we have advocated in this review and described in Box 1. It is nearly 15 years since Jean Bullier posed the question “What is fed back?” (Bullier, 2006). The multiplicity of FB pathways and the complexity of their proposed functions were deep issues that he felt needed to be addressed. In the last 14 years there has been a spectacular development of three classes of techniques that now allow us to address Jean Bullier’s question. Firstly, optogenetics holographic stimulation makes it possible to address causation (Carrillo-Reid et al., 2019; Marshel et al., 2019), thereby addressing how activation of a given FB pathway influences a particular cognitive task. Secondly, viral tracing allows cell-type and laminar-constrained connectivity (El-Shamayleh et al., 2016; Nassi et al., 2015; Siu et al., 2020) making it possible to resolve the dual counterstream architecture. Thirdly, high-resolution fMRI allows laminar and columnar resolution imaging in the human brain e.g. (Kemper et al., 2018); furthermore, recent advances in MR methods and data analysis enable investigation of directional laminar connectivity in hierarchical brain networks (Huber et al., 2020). The key feature of future investigation will be to examine FB modulation of cognitive tasks in animal models that can, in parallel, be applied in humans where perceptual consequences can be reported (Kok et al., 2016; Schneider et al., 2019). These combined approaches will address the complexity of the interaction of descending generative networks with the local microcircuit. Interpreting the data coming out of these experiment in terms of cortical dynamics will require computational modeling of the interactions of these top-down effects with the canonical microcircuit using approaches developed by Maass (Haeusler and Maass, 2007).

The interactions of descending with ascending pathways required by predictive processing theory will be largely in supragranular cortical layers, which have been shown to be a major target for human evolutionary adaptation (Heide et al., 2020; Won et al., 2019). The evolutionary expansion of the supragranular layers is accompanied by an increase in heterogeneity of glutamergic cell-types in terms of morphology, electrophysiology and gene expression going from rodent to human (Berg et al., 2020) in turn supporting a complexification of the circuits in these layers (Hodge et al., 2019). The amplification and diversification of supragranular intratelencephalic-projecting neurons in primates suggest that the investigation of the biology of the generative networks advocated here may well exploit reductionist approaches in the rodent model, but will need nonetheless to be studied with a particular emphasis on human and non-human primates.

Finally, the observation that top-down networks generate sensory-like activity in low levels of the cortical hierarchy in the *absence* of sensory input raises interesting issues of representation in the brain. According to predictive coding theory, sensory perception requires predicting the present. These top-down generative networks can be important for predicting or simulating the future (Grezes and Decety, 2001)<u>, in interplay with the hippocampus (Buckner, 2010; Kay et al., 2020),</u> as well as revisiting the past allowing for instance the minds eye to scrutinize detail in visual images (Kosslyn, 1994). In this way the top-down generative networks could have much to do with what we call thinking. In addition these networks would support an integral aspect of human consciousness, namely the seamless and continuous navigation of the mental narrative of self across past, present and future (Jaynes, 1976; Nørretranders, 1991). Importantly, the observation that these networks are active in the NHP <u>(van Kerkoerle et al., 2017), i</u>ndicates that we can now plan experimental investigation of their integrative neurobiology at the cellular level.

## Acknowledgments

We would like to thank Kevan Martin and Rodney Douglas for numerous and often passionate discussions on some of the central issues discussed here and for their thoughtful comments on the text, as well as Stewart Ship for his deep insight. Funding gratefully acknowledged by XJW (ONR Grant N00014-17-1-2041, US National Institutes of Health NIH grant 062349, Simons Collaboration on the Global Brain program grant 543057SPI); KK (DUAL_STREAM ANR-19-CE37-0025); MV (ERC Starting Grant SPATEMP, BMF BINDA-031L0167); HK (LABEX CORTEX ANR-11-LABX-0042; Université de Lyon ANR-11-IDEX-0007), A2P2MC ANR-17-NEUC-0004, CORTICITY ANR-17-HBPR-0003, SCUSI, Région Auvergne-Rhône-Alpes 1700933701, FRC APE13 “Rotary-Espoir en Tête”.

## Author contribution

Data acquisition JV, LM, HK; Analysis of data JV, LM, KK, HK; Statistical modeling of hierarchy KK; All authors conjointly wrote the first draft and revised and edited the completed document; Proposed the study MV, HK

